# Light of both high and low melanopic illuminance improves alertness and attention during daytime

**DOI:** 10.1101/2025.01.15.633100

**Authors:** Louise Bruland Bjerrum, Endre Visted, Inger Hilde Nordhus, Berge Osnes, Bjørn Bjorvatn, Oda Bugge Kambestad, Malika Elise Hansen, Lin Sørensen, Elisabeth Flo-Groeneboom

**Affiliations:** Department of Clinical Psychology, University of Bergen, Norway; Department of Global Public Health and Primary Care, University of Bergen, Norway; Norwegian Competence Center for Sleep Disorders, Haukeland University Hospital, Norway; Department of Biological and Medical Psychology, University of Bergen, Norway

## Abstract

Light exposure during the day exerts acute effects on attention, such as how alert and ready a person is for solving problems and goal-oriented behavior. However, to increase the understanding of how different light conditions during daytime affect our attention, there is a need for more studies. The current study, using a sample of healthy, young adults (*N* = 39; 21.7±2.6 y, 61.5% female), tested the acute effects of morning exposure (09:00-11:00) to four artificial lights on attention and alertness with the Psychomotor Vigilance Task (PVT). The PVT was administered three times in short-wavelength “blue” light [SWL, high melanopic illuminance], long-wavelength “red” light [LWL, low melanopic illuminance], and bright white light [BWL] of high correlated color temperature, and white dim light [DL] as control condition. PVT measures included fluctuations in attention, quantified as intra-individual Reaction Time (RT) variability and the number of lapses; mean RT; and optimum response capability (10% fastest RTs). Compared to DL, participants had significantly fewer lapses and faster mean RT during SWL and LWL, and they enhanced their optimum responses in LWL. There were no significant effects of BWL, and light did not affect RT variability. Surprisingly, SWL was not superior to LWL. Hence, our results suggest that light of both high and low melanopic illuminance can improve alertness and attention in the morning.

## Introduction

Impaired alertness is a common problem in everyday life and can arise from inadequate sleep related to work schedules and various societal/domestic demands. Irrespective of the cause, low alertness levels can be a risk factor for accidents and reduced performance in occupational and operational settings, especially during night shifts^1–7^. Exposure to light has been tested as an intervention to optimize alertness during night shifts, however, most of these interventions have involved exposure to white light^8–10^. It is therefore of interest to better understand how different light conditions affect aspects of alertness, also during the day, to enhance cognitive performance.

Light is involved in a range of so-called Non-Image-Forming (NIF) responses, including acute effects on alertness^11,12^, melatonin suppression^13–15^, pupil constriction^16,17^, and synchronization of the “master” circadian clock in the brain to the environmental light-dark cycle^18^. NIF responses are mainly driven by intrinsically photosensitive Retinal Ganglion Cells (ipRGCs), a small subset of photoreceptors in the retina^19^. ipRGCs contain the photopigment melanopsin that is maximally sensitive to short-wavelength light (SWL) with a peak at ∼ 480 nm^20,21^. SWL with a narrow bandwidth (that is, with a narrow peak in some part of the visible electromagnetic spectrum) will appear blue to the human eye. Similarly, medium-wavelength light (MWL, with a peak spectral wavelength at ∼ 550 nm) will appear green, and long-wavelength light (LWL, with a peak spectral wavelength at ∼ 630 nm) will appear red. Polychromatic white light will also activate melanopsin, especially if the white light is bright enough, and/or “cool”. “Cool” white light is often enriched in shorter wavelengths, resulting in high Correlated Color Temperature (CCT, measured in kelvin (K)), whereas “warm” light is enriched in longer wavelengths, resulting in lower CCT. Light with high proportions of short wavelengths will typically also have a high melanopic-Equivalent Daylight Illuminance (m-EDI). This metric has been proposed as the preferred metric (over the “traditional” photopic illuminance metric) when quantifying the biological effects of light^22,23^. Given the role of light in various NIF responses it is crucial to understand how different light conditions influence attention and alertness. Alertness, a subfunction of attention, follows a circadian (∼24-hour) rhythm, with low alertness levels immediately upon awakening, followed by increased and rather stable levels throughout the day and subsequent lower levels in the evening and at night^24^. Being alert and attentive is associated with the functioning of the alerting network, one of the three main attention networks of the human brain^25^. Adequate alertness levels throughout the day enable high sensitivity to incoming stimuli and facilitate more complex, cognitive tasks^26^. Studies have shown that exposure to SWL, bright white light (BWL), and short-wavelength-enriched white light during the day can increase neurobehavioral alertness^12,27–30^ and reduce brain activity in frequency bands (electroencephalography (EEG)) associated with sleepiness/drowsiness^16,27,31,32^. However, other studies report no significant effects^11,29,33,34^. Furthermore, studies show that exposure to LWL can also increase alertness as indicated by effects on EEG markers^11,31,32^. Importantly, though, acute alerting effects of light can be influenced and confounded by a range of factors, such as chronotype (inter-individual differences in preferred sleep and wake timing, e.g., whether one is a “morning lark” or an “evening owl”)^35^ and various sleep-related variables. Poor sleep quality and sleep restriction (curtailed sleep duration) have been linked to alertness deficits^36–39^, however, light exposure has the potential to counteract such deficits^40–42^. Furthermore, seasonal changes in ambient light levels can also affect NIF responses to light, probably due to changes in light sensitivity^43,44^. Therefore, it is important to take these factors into consideration when assessing the alerting effects of light.

The Psychomotor Vigilance Task (PVT) is a high signal-load reaction time (RT) task widely used to assess alertness in laboratory contexts. It has minor learning effects, making it ideal for repeated measures^45^. The most commonly reported PVT metrics are mean reaction time (RT) and the number of lapses in attention (often quantified as RTs ≥ 500 ms). A less commonly reported PVT metric is the 10% fastest RTs, reflecting the optimal responses on the task^46^. Studies using the PVT to assess alertness in different light conditions have reported faster mean RT and fewer lapses during SWL (“blue”)^27^, bright white light^12,28,47^ and white light of high m-EDI^29^. However, null findings have been reported as well^34,48–53^.

Additionally, most studies report no significant effects of white light on the 10% fastest RTs^12,28,30,48–50,52,54–56^. When it comes to the effects of LWL on PVT measures or other sustained attention tasks, studies are scarce and the results are inconclusive^32,53^. Also, it remains unknown whether the 10% fastest RTs in particular can be modulated by exposure to narrow bandwidth lights during the day. Finally, relatively few studies have assessed light’s impact on fatigue over time in testing, a factor that increases RTs across PVT sessions^12,47,48,51^.

In sum, research on the acute effects of light during daytime on neurobehavioral alertness is inconclusive. Importantly, previous literature on the NIF acute effects of light on RT-based alertness has focused on central tendency measures, reflecting average performance across individuals. To our knowledge, it remains to be tested whether different types of artificial light can exert acute effects on alertness and attention measured at an intra-individual level in terms of RT variability (RTV). As RTV on the PVT is underreported in the literature, it is of interest to include this outcome to get a more nuanced picture of how light acutely impacts attentional fluctuations. Also, there is limited knowledge on how chronotype and season might affect neurobehavioral performance during light exposure. The overall aim of the present study was therefore to assess the impact of daytime exposure to different artificial light conditions on neurobehavioral alertness in healthy, young adults. By assessing not only the acute effects of different light conditions on central tendency measures, this study also tests the acute effects of light on intra-individual RTV, thereby filling a gap in the literature. Hypotheses were pre-registered at OSF (https://osf.io/m7gfh). Based on previous research and theory on the NIF acute effects of light, we hypothesized that:

- Short-wavelength (“blue”) light (SWL), long-wavelength (“red”) light (LWL) and bright white light (BWL), compared to dim light (DL), will reduce fluctuations in attention (quantified as intra-individual RT variability and the number of lapses) and overall mean RT on the PVT.
- SWL, LWL, and BWL, compared to DL, will not affect optimum response capability (quantified as the mean of the 10% fastest RTs).

As the differential effects between SWL, LWL, and BWL on alertness and attention are inconclusive in the literature^57^, we tested the following explorative hypothesis:

- SWL will have a superior effect on intra-individual RTV, lapses and mean RT, compared to LWL, and BWL will have a superior effect on intra-individual RTV, lapses, and mean RT, compared to LWL.

Last, we also hypothesized, exploratively, that:

- Participants tested in fall and winter will show stronger responses to light than participants tested in spring and summer.
- Individuals with a ‘morning’ chronotype or ‘intermediate’ chronotype will perform better than ‘evening’ chronotypes, and ‘evening’ chronotypes will benefit more from light with high m-EDI compared to DL, relative to ‘morning’ and ‘intermediate’ chronotypes.

## Methods

### Study design and setting

This study employed a within-subjects repeated-measures experimental design to test the acute effects of daytime exposure to different artificial light conditions on alertness and attention in healthy, young adults. Data were collected from November 2021 to October 2023, at a high latitude (60.39° N) in a 23 m^2^ ventilated laboratory with no windows, white painted walls, and 20 60×60 ceiling-mounted Light Emitting Diode (LED) luminaires (Modul R 600 LED CCT/RGB MP; Glamox AS, Norway). See Fig. S1 for an overview of the laboratory setup. The laboratory had nine workstations in total. Each workstation measured approximately 1.15 meters in width and was equipped with a desk, chair, a monitor, keyboard, and a computer mouse. Two luminaires were situated directly above each workstation, providing uniform illumination. Workstations were physically separated by gray partition walls. All monitors had an overlay “orange” screen foil blocking wavelengths < 520 nm (METOLIGHT®SFG10, Asmetec, Germany).

### Participants

Healthy, young adults (≤ 40 years old) were recruited from lectures at the University of Bergen (Norway) and via online/on-campus posters. Based on a screening procedure, participants were included if they had normal or corrected-to-normal vision. Exclusion criteria included colorblindness (as indicated with Isihara’s Test for Color Deficiency), recent transmeridian travel (crossing three time zones within the last three weeks), a severe psychiatric disorder (screened by the SCL-90 and MINI module C for mania), or a severe medical disorder.

### Procedures

Participants underwent one week “at-home-baseline” with usual routines, followed by four morning lab-sessions from 08:00 to 11:00 on separate days across two weeks. For each of the two weeks, participants were scheduled to participate either on Monday and Wednesday or on Tuesday and Thursday. One light condition was presented per session in a randomized block-stratified counterbalanced order for each participant to partly balance out learning effects and effects related to differences in circadian phase. Throughout the three weeks of study participation, participants were instructed to maintain a regular sleep-wake schedule (including weekends), however, they were not required to spend a predefined number of hours in bed each night. Sleep was tracked with diaries and Actiwatch Spectrum Plus devices (Philips Respironics, Inc., United States) worn on participants’ non-dominant wrists. On the morning of scheduled test sessions, participants were instructed to wear short-wavelength light-blocking glasses (Uvex, Honeywell International, Inc., United States) immediately upon awakening to minimize circadian phase shifts, which are more pronounced with exposure to light spectra with high proportions of short wavelengths^13,58,59^. Habitual morning caffeine consumption was allowed before entering the laboratory on the days of scheduled test sessions, however, no caffeine consumption nor nicotine use was permitted in the laboratory. Participants arrived at the laboratory at 08:00 and were maintained in warm-white dim light (DL, m-EDI = 4.8±6.2 lux (lx)) for one hour. Electrocardiogram electrodes were applied, and resting heart-rate variability (HRV) was sampled at ∼08:40. From ∼08:50, the assigned experimental light condition increased gradually in illuminance over 10 minutes to limit eye strain. Then, participants underwent a 30-minute light adaptation in the assigned light condition. During ECG setup and light adaptation, participants sat at their workstations, were allowed to converse and received a small snack. Water was available during the entire test session. Pupils were not pharmacologically dilated prior to light exposure. After light adaptation, a second 10-minute resting HRV was sampled, followed by the completion of neurobehavioral tasks for approx. 1.5 hours during exposure to the assigned light condition (see Fig. 1).

**Figure 1.**
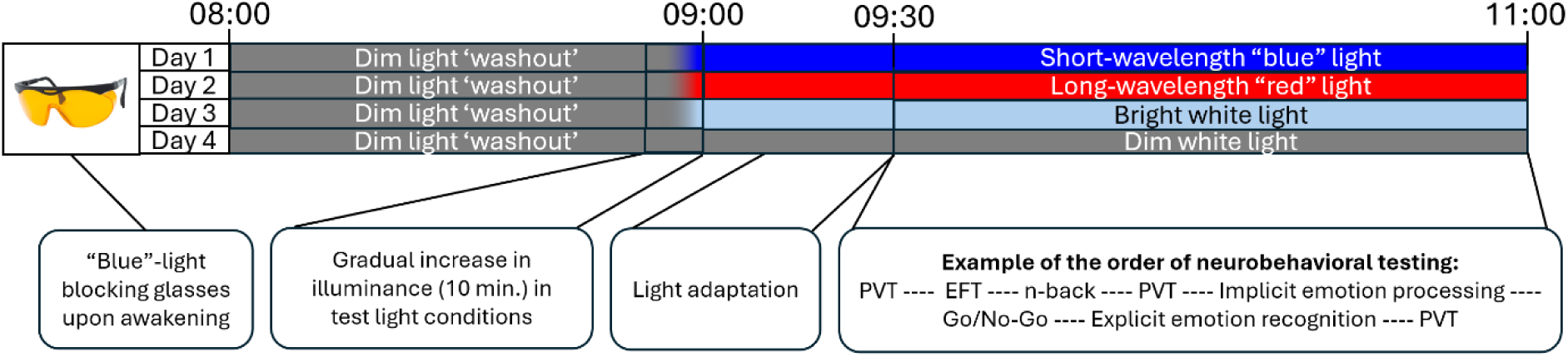
Overview of experimental design. From 09:30 to approx. 11:00, participants performed a series of neurobehavioral tests assessing alertness (Psychomotor Vigilance Task (PVT)), cognitive control (Eriksen Flanker Task (EFT)), working memory (n-back), and aspects of emotion processing (implicit and explicit emotion recognition and a Go/No-Go task with emotional stimuli). Results from the EFT, n-back. and emotion tasks will be reported in future publications. The presentation order of light conditions and the order of cognitive vs. emotion-related tests were randomized and cross-balanced within groups of participants, however, not for the PVT, as this task was administered in three blocks at fixed intervals throughout light exposure (at ∼09:40, ∼10:15, and ∼10:50).

Experimental light conditions were: narrow bandwidth SWL (“blue”, ʎ_max_ = 455 nm, m-EDI = 1441.8±35.2 lx), narrow bandwidth LWL (“red”, ʎ_max_ = 625 nm, m-EDI = 3.8±1.3 lx), and SWL-enriched BWL (m-EDI = 1155.8±17.6 lx, CCT = 8035.7±42.8 K). Hence, SWL and BWL were both of relatively high m-EDI, whereas the LWL condition had a low m-EDI. SWL and LWL were photon-matched (irradiance = log 18.8/m^2^/s). Effects of SWL, LW, and BWL were contrasted against a control condition of white DL (m-EDI = 9.7±6.4 lx). Light quantities were calculated using the open-access- and source web platform *luox*^22,60^. Detailed light quantities, spectral power distribution curves, and spectral power raw data can be accessed in Table S1, Fig. S2, and Table S2, respectively. SWL, LWL, and BWL were delivered through the ceiling-mounted luminaires. In the DL control condition, all LED luminaires were off, and only computer screen light was present in the room. During the one-hour exposure to warm-white DL from 08:00 to 09:00, preceding light adaptation, DL was delivered through six portable Philips Hue Go lamps positioned behind each monitor facing upward. At each workstation, light was measured with a calibrated spectroradiometer (GL Spectis 1.0 T Flicker, GL Optic, Poland) at eye level (vertically, 120 cm above the floor), ∼90 cm horizontally from the computer screen in the direction of gaze (corresponding to the approximate seating of participants) with the screens turned on. The light measurements were performed approx. 30 minutes after the onset of the LED luminaires to allow the luminaires to adjust in temperature.

### Ethics

Participants provided written informed consent before participation and received 1000 NOK (∼ 94 USD) after completing the study. Ethics approval was provided by the Regional Committees for Medical and Health Research Ethics. All procedures contributing to this work comply with the Declaration of Helsinki.

### Measures

#### Alertness

Alertness was assessed three times in each of the four light conditions (block 1-3, see Fig. 1) with a five-minute computerized version of the PVT^45,61^ administered through E-Prime 3.0^62^. Participants monitored a red rectangular box on a white background. When a counting timer (stimulus, in black) appeared inside the box and started counting, participants were instructed to press the space bar as quickly as possible. The stimulus was presented at random intervals ranging from 2 to 7 s, and 1-s feedback on RT was provided after each trial. RTs < 100 ms were considered false starts. Five practice trials were presented before each test. The mean number of trials in each PVT administration was 57. Outcome measures reported here are the number of lapses in attention (quantified as RTs ≥ 500 ms), overall mean RT, intra-individual RTV (quantified as the standard deviation (SD) in RTs^63^), and optimum response capability (quantified as the mean of the 10% fastest RTs).

#### Covariates

Chronotype was assessed with a reduced version of the Horne & Östberg Morningness-Eveningness Questionnaire (rMEQ)^64^. The rMEQ has five questions probing individuals’ circadian preferences, yielding a total score ranging from 4 to 25. A score of 4-7 = ‘Definitely Evening Type’, 8-11 = ‘Moderately Evening Type’, 12-17 = ‘Intermediate Type’, 18-21 = ‘Moderately Morning Type’, and 22-25 = ‘Definitely Morning Type’. ‘Definitely evening type’ and ‘definitely morning type’ is often referred to as “extreme” chronotypes.

Season was not systematically manipulated between nor within subjects (each subject underwent laboratory testing in one season only, across a study period of three weeks) but was entered as a covariate in statistical analyses. As there is substantial variation in annual photoperiod length at northern latitudes^65^, all four seasons were included as categorical levels. December-February represented ‘winter’, March-May represented ‘spring’, June-August represented ‘summer’, and September-November represented ‘fall’.

Sleep efficiency the night preceding each test session was derived from wrist actigraphy data, recorded in epochs of 1-minute duration. Threshold sensitivity was set to ‘medium’ (40 counts per minute) (Actiware version 6.0.9, Philips Respironics, Inc., United States). Rest intervals were scored based on i) the participants pushing an event button together with ii) raw data, and iii) information from sleep diaries. When the event button was not pushed, information from the latter two data sources was used. For each rest interval, sleep efficiency was calculated as: (total sleep time/(interval duration – total invalid time))*100.

### Statistical analyses

The analysis plan was pre-registered at OSF, together with the hypotheses (https://osf.io/m7gfh). Missing data (< 10%) were imputed with a Random Forest algorithm in the R package *mice*^66^. All analyses were performed in R version 4.4.1^67^ using the packages *glmmTMB*^68^, *lme4*^69^, *lmerTest*^70^, *merDeriv*^71^, *DHARMa*^72^*, performance*^73^, *parameters*^74^*, correlation*^75^, *emmeans*^76^, and *effectsize*^77^.

Data from the five practice trials on the PVT were excluded from analyses. First, a set of initial exploratory analyses was undertaken to rule out the effect of variables not directly related to light that can confound the results. As caffeine consumption can affect attention^78^, each PVT outcome was analyzed with either a generalized linear mixed-effects model (GLMM) or a linear mixed-effects model (LMM) depending on the distribution of residuals, controlling for whether participants habitually consumed caffeine or not (“Yes”, “No”, respectively). Subject was modeled with random intercept. Also, even though learning effects related to repeated test administrations across the four light conditions were partly balanced out by randomizing the presentation order of light conditions for each participant, learning effects on PVT measures were analyzed with GLMMs or LMMs by including session number (reflecting the presentation order for each light condition) as predictor, a random intercept for subject and a complex random intercept (CRI)^79^ for the random effect of session number.

#### Primary analyses

The number of lapses did not exhibit significant zero-inflation nor overdispersion and was therefore analyzed with a GLMM using a generalized Poisson distribution. RTV was also analyzed with a GLMM due to right-skewness in the residuals. Mean RT for all trials and the mean of the 10% fastest RTs were each analyzed with a LMM using Restricted Maximum Likelihood (REML). As the residuals of RTV, mean RT, and mean 10% fastest RTs deviated slightly from a normal distribution, variables were log-transformed prior to analysis to increase normality and stabilize variance. In all analyses on PVT variables, light*block, chronotype, season, and sleep efficiency were modeled as fixed effects. Subject was modeled with random intercept, and the two other within-subjects factors of light and block were modeled with CRIs^79^. In the analysis of RTV, mean RT was added as an additional predictor because the mean and SD of RTs are often correlated^80^. Primary analyses included both interaction models (light*block) and main effects models (light and block modeled as separate fixed effects).

#### Secondary exploratory analyses

Exploratory analyses included models with the interaction term light*chronotype in one model and light*season in another model, controlling for the random and remaining fixed effects specified above, including block as main effect. We also explored sex differences on all PVT measures, as there are some indications that sex influences acute alerting responses^81^.

#### Statistical inference and approximation of effect sizes

Alpha level was set to .05 (two-tailed) in all analyses. Significant main and/or interaction effects were followed up with Tukey’s Honest Significant Difference (HSD) test. Partial *r* (*r*_p_) (GLMM estimates) and Cohen’s partial *f*^2^ (LMM estimates) were calculated as theoretical effect sizes^82^ and were approximated using the test-statistic approximation method^77^. Small effect sizes corresponding to a *r*_p_ < .3 and *f*^2^ ≥ .02, medium effect sizes equal to *r*_p_ < .5 and *f*^2^ ≥ .15, and large effect sizes correspond to *r*_p_ ≥ .5 and *f*^2^ ≥ .35^83^.

## Results

40 healthy, young adults participated in the study. 42 out of a total of 480 observations were excluded from analyses due to measurement errors, resulting in a sample of 39 participants (mean age: 21.7, SD = 2.6, range: 18-28) and 438 observations (Table 1). Mean rMEQ score was 13.8 (SD = 3.3), corresponding to an ‘intermediate’ chronotype. On average, participants went to bed at 23:47 (SD = 01:14) and woke up at 06:41 (SD = 32 minutes), resulting in an average sleep duration of 372 minutes (∼ 6.2 hours, SD = 69.8 minutes). Upon arrival at the laboratory at 08:00, participants had been awake for an average of one hour and 18 minutes (SD = 32.4 minutes). On the PVT, irrespective of light manipulation and measurement block, participants’ median number of lapses were 1 (range: 0-16), mean overall RT was 308.40 ms (SD = 40.50 ms), mean intra-individual RTV was 75.31 ms (SD = 71.27 ms), and the mean of the 10% fastest RTs was 242.96 ms (SD = 21.79 ms).

**Table 1.** Descriptive statistics of study sample and laboratory characteristics. Chronotype was assessed with a reduced version of the Morningness-Eveningness Questionnaire. There were no extreme chronotypes. Sleep efficiency was derived from wrist actigraphy. Caffeinated beverages represent the median number of caffeinated beverages (coffee, caffeinated tea, energy drink) habitually consumed on a normal day. *Kruskal-Wallis Rank Sum Tests did not reveal any significant differences in temperature (*p* = .251) nor air humidity (*p* = .539) between light conditions.

|  |  |
| --- | --- |
| Total N (females) | 39 (24) |
| Age [Mean±SD] | 21.7±2.6 |
| Caffeinated beverages [Median (range)] | 1 (0-5) |
| Sleep efficiency [Mean±SD, %] | 83.5±8.9 |
| **Chronotype N (%)** |  |
| Definitely morningness | 0 (0) |
| Moderately morningness | 7 (17.9) |
| Intermediate type | 22 (56.4) |
| Moderately eveningness | 10 (25.6) |
| Definitely eveningness | 0 (0) |
| **Season N (%)** |  |
| Spring | 10 (25.6) |
| Summer | 5 (12.8) |
| Fall | 18 (46.2) |
| Winter | 6 (15.4) |
| **Laboratory conditions\*** |  |
| Temperature [Mean±SD, °C] | 20±1.2 |
| Humidity [Mean±SD, %] | 39.8±13.2 |

Approx. 64% of participants habitually consumed one or more caffeinated beverages on a normal day, however, exploratory mixed-model analyses confirmed that caffeine consumption did not significantly affect performance. Neither were there any clear indications of learning effects on PVT measures, as indicated by more lapses and slower RTs across some sessions rather than improvements in performance, which would have been expected if there were prominent learning effects. Additional exploratory analyses showed no significant sex differences in performance.

### Acute alerting effects of light

Line plots of the progress in RTs from trial 1 to 54 in each light condition, stratified by measurement block, are provided in Fig. S3, Fig. S4, and Fig. S5. As the initial adjusted models with the interaction term light*block revealed significant simple effects of light and/or block but no significant interactions, light and block were modeled as separate fixed effects in subsequent main effects models. Results from these models are described below. Tables with model estimates, effect sizes, and explained variance statistics (Nakagawa’s *R*^2^ and the intra-class correlation coefficient (ICC)) can be accessed in Table S3, Table S4, Table S5, and Table S6. On all PVT measures except for RTV, participants responded slower over time (main effects of measurement blocks; *p*’s < .001, see Fig. S6a, Fig. S7, Fig. S8, and Fig. S9a).

#### Lapses

The analysis of the number of lapses showed significant main effects of light (SWL: *p* = .004, *r*_p_ = -0.42; LWL: *p* = .002, *r*_p_ = -0.44; BWL: *p* = .044, *r*_p_ = -0.31, Table S3). Post-hoc analyses showed that participants had significantly fewer lapses in SWL (*p* = .023, *EMM* = 0.89[0.63-1.27], *SE* = 1.20) and LWL (*p* = .010*, EMM* = 0.85[0.59-1.22], *SE* = 1.20), compared to DL (*EMM* = 1.48[1.05-2.10], *SE* = 1.20) (Fig. 2a). There was also a significant main effect of season (spring vs. fall: *p* = .027, *r*_p_ = 0.33, Fig. S6b), but the effect was no longer significant after correcting for multiple comparisons. Also, the model with the interaction term light*season revealed a significant interaction between SWL and winter (*p* = .025), but this effect was also no longer significant after correcting for multiple comparisons. There were no significant light*chronotype interactions.

**Figure 2.**
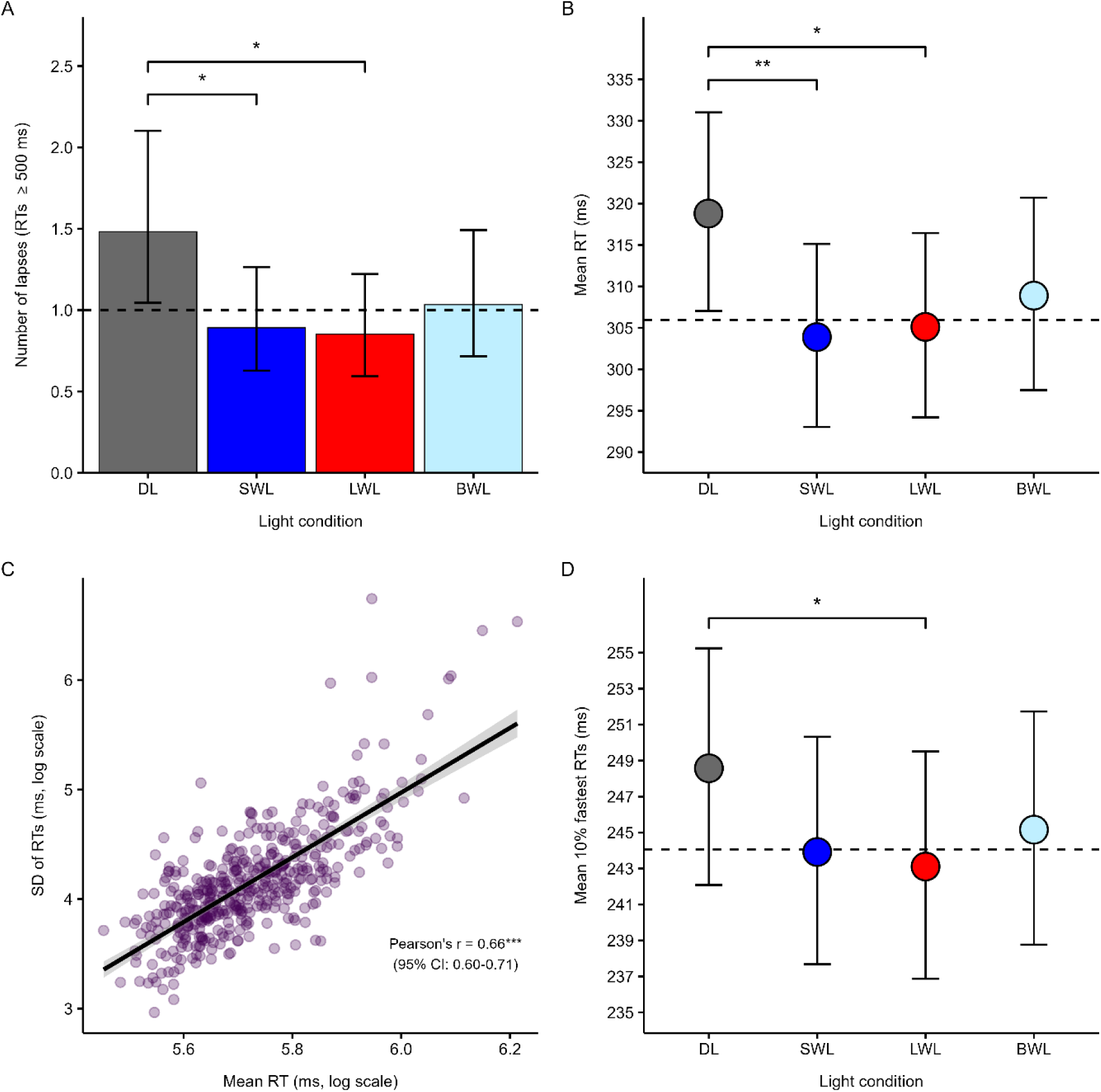
Y-axes in panel A, B, and D show model-based estimated marginal means (EMMs) for the main effect of light, with error bars representing the 95% confidence interval (CI). EMMs and CIs were back-transformed from log to original scale for visualization purposes. (A) EMMs for the main effect of light on the number of lapses (RTs ≥ 500 ms). Dashed line represents the median number of lapses across conditions. (B) EMMs for the main effect of light on mean RT (in ms). The y-axis limit has been adjusted so that group differences appear more clearly. Dashed line represents mean estimated RT across SWL, LWL, and BWL. (C) Scatter plot of the relationship between RTV, quantified as the standard deviations in RTs (in ms), and mean RT (ms). There was a strong correlation between the SD and the mean in RTs (Pearson’s partial *r* = .66). (D) EMMs for the main effect of light on mean 10% fastest RTs. The y-axis limit has been adjusted so that group differences appear more clearly. Dashed line represents the estimated mean 10% fastest RT across SWL, LWL, and BWL. BWL = Bright White Light. DL = Dim Light. LWL = Long-Wavelength Light. ms = milliseconds. RTs = Reaction Times. SWL = Short-Wavelength Light. \*\*\**p* < .001. \*\**p* (corrected) < .01. \**p* (corrected) < .05.

#### Mean RT

The main effects model showed significant main effects of light (SWL: *p* = .001, *f*^2^ = 0.11; LWL: *p* = .004 *f*^2^ = 0.09; BWL: *p* = .038 *f*^2^ = 0.04, Table S4). Post-hoc analyses revealed that participants’ mean RT was faster in SWL (*p* = .008, *EMM* = 303.89[293.05-315.14], *SE* = 1.02) and LWL (*p* = .018, *EMM* = 305.12[294.20-316.45], *SE* = 1.02) compared to DL (*EMM* =

318.81[307.05-331.02], *SE* = 1.02, Fig. 2b). Exploratory analyses showed no significant interactions between light and season nor between light and chronotype.

#### RTV

The main effects model showed a significant main effect of mean RT (*p* < .001, *r*_p_ = 0.943, in that variability in RT increased with longer RTs (Pearsons’s partial *r* = .66 [95% CI: .60-.71], *p* < .001, Fig. 2c). There were also significant main effects of measurement blocks (block 1: *p* < .001, *r*_p_ = -0.51; block 2: *p* < .001, *r*_p_ = -0.61, Table S5). Post-hoc analyses showed that participants expressed lower variability in block 2 (*p* < .001, *EMM* = 60.61[55.76-65.88], *SE* = 1.04) and 3 (*p* < .0001, *EMM* = 58.03[53.30-63.18], *SE* = 1.04), relative to block 1 (*EMM* = 69.70[64.04-75.87], *SE* = 1.04), see Fig. S8. There were no significant main effects of light, season or chronotype, and no significant interactions.

#### Mean of the 10% fastest RTs

The main effects model on the mean of the 10% fastest RTs showed significant main effects of light (SWL: *p* = .017, *f*^2^ = 0.06; LWL: *p* = .005, *f*^2^ = 0.08), chronotype (eveningness: *p* = .019, *f*^2^ = 0.03; morningness: *p* = .023, *f*^2^ = 0.02), and season (summer: *p* = .018, *f*^2^ = 0.16), see Table S6. Post-hoc analyses showed that participants responded faster in LWL (*p* = .030, *EMM* = 243.11[236.88-249.51], *SE* = 1.01) compared to DL (*EMM* = 248.57[242.08-255.24], *SE* = 1.01, Fig. 2d). Post-hoc analyses on the main effect of season showed that participants tested in winter responded faster (*p* = .048, *EMM* = 234.68[222.32-247.73], *SE* = 1.03) than participants tested in summer (*EMM* = 261.25[246.19-277.23], *SE* = 1.03, Fig. S9b). Post-hoc analyses of the main effects of chronotype did not remain significant after correction for multiple comparisons, and the interaction models of light*season and light*chronotype showed no significant interactions.

### Additional Results

#### Sensitivity Analyses

Sensitivity analyses on a dataset with complete-cases only (listwise deletion) showed a similar overall pattern of results (see Supplementary Materials S4) as in analyses on imputed data, although *p*-values were generally lower and standard errors were smaller, as expected in complete-case analyses when missing data is not missing completely at random (MCAR)^84,85^. The overall pattern of results was similar to the results from analyses on imputed data (see Fig. S10, Fig. S12, Fig. S13, and Fig. S14. The only results in complete-case analyses that deviated from results obtained from analyses of imputed data were a significant interaction between light and chronotype on lapses (DL eveningness*SWL intermediate type, *p*_corrected_ = .048, Fig. S11), and a significant main effect of light on RTV (larger variability in LWL compared to DL; *p_corrected_* = .031, Fig. S13a). However, these results should be interpreted with caution, due to the mentioned biases often introduced in estimates when the missing data is not MCAR.

#### Crude Models

In addition to the primary main-effects models on imputed data, we also constructed “crude” main-effects models unadjusted for the influence of chronotype, season, and sleep efficiency, with light and block included as the only fixed effects, a random intercept for subject, and a CRI for light and block. The main effects of light and block in these analyses were similar to the results from the primary main-effects model analyses reported below. Model estimates from “crude” models are provided in Table S7, Table S8, Table S9, and Table S10.

## Discussion

This crossover study tested whether a 2-hour (09:00-11:00) exposure to artificial light of high and low melanopic-equivalent daylight illuminance (m-EDI) could improve alertness and attention in healthy, young adults. Beyond standard central tendency measures from the PVT (lapses, mean RT, and mean 10% fastest RTs), we report – to our knowledge – the first test of light’s acute effect on intra-individual RT variability (RTV). Results partly supported our main hypotheses; that narrow bandwidth SWL and LWL, relative to white DL, reduced lapses and shortened mean RT. LWL also improved the 10% fastest RTs. Light did not exert acute effects on RTV. Surprisingly, there were no significant effects of BWL on either of the PVT measures, despite the fact that the BWL condition was also of relatively high m-EDI as well as high CCT. And, contrary to one of our explorative hypotheses, we did not find evidence supporting that SWL was superior to LWL in inducing alerting effects. Hence, irrespective of m-EDI, both SWL and LWL improved alertness, vigilance, and optimal responding in a sample of healthy, young adults in the morning.

Relatively few daytime studies have tested the acute effects of narrow bandwidth SWL on alertness, and none of these have reported melanopic light quantities^27,31,34,53^. Additionally, substantial heterogeneity in light protocols among studies complicates comparisons. Nevertheless, our finding that SWL, relative to white DL, reduced the number of lapses and mean RT aligns with Rahman et al.^86^, though their control condition was medium-wavelength “green” light and exposure duration was much longer (6.5 hours). Laszewska et al.^31^ similarly showed that 40 minutes of SWL of low photopic illuminance in the early afternoon reduced EEG power in frequency bands associated with drowsiness/sleepiness, whereas van der Meijden et al.^53^ showed that intermittent SWL (5-minute light blocks separated by blocks of darkness) did not affect mean RT on the PVT, suggesting that continuous light exposure may be critical for SWL to induce alerting responses. A plausible mechanism underlying the alerting potential of SWL during daytime is the light-induced engagement of brain areas in the alerting network – including the dorsolateral prefrontal cortex (DLPFC), the anterior cingulate cortex (ACC), and the Locus Coeruleus (LC) in the brain stem^87,88^. Moreover, stable PVT performance correlates with greater functional connectivity between the DLPFC and other cortical areas^34^. However, to date, only one study has tested the impact of SWL on the efficiency of the alerting network, as measured with the Attention Network Test (ANT)^89^. This study reported no significant effect of SWL administered in the afternoon^90^, however, the light exposure duration was rather brief (5 minutes). Meanwhile, an fMRI study by Vandewalle et al.^87^ showed that 50 seconds of SWL administered approx. 4 hours after habitual wakeup time was sufficient to induce activations in brain areas associated with the alerting network. Nevertheless, it currently remains unknown whether longer exposures to narrow bandwidth SWL exert a different effect on the alerting network, and which specific light characteristics, such as m-EDI and time-of-day, are required to induce these effects.

Beyond acute alerting effects of SWL on lapses and mean RT, we also observed similar effects during exposure to LWL. Again, as for the literature covering acute effects of SWL on neurobehavioral alertness, previous studies testing the effects of narrow bandwidth LWL are rather scarce, and these report no significant effects of LWL^32,53^. Although the light exposure duration and control light condition in the study by Sahin et al.^32^ were comparable to our study, a direct comparison of results is still somewhat complicated, as the study used a task probing alertness over a longer time period (i.e., sustained attention^91^) and the brightness of LWL was much lower (213 vs. ∼ 560 photopic lx in our study). The absence of a significant effect of LWL on PVT mean RT reported in one of the experiments in van der Meijden et al.^53^ might be due to the intermittent light exposure pattern, which differs from our continuous exposure. Also, the illuminance of LWL in van der Meijden et al.^53^ was only 40 photopic lx, which might not be sufficient to induce alerting responses during the day. When it comes to other PVT measures, our study is the first to test the acute daytime effects of LWL on lapses and the most optimal responses (mean 10% fastest RTs). Hence, future studies are needed to confirm our results showing a reduction in attentional lapses and enhanced optimum responses during exposure to bright LWL. At a neurophysiological level, studies have shown acute alerting effects on EEG markers of alertness during LWL exposure ranging from 40 to 110 minutes in duration^11,31,32^. Additionally, it has been shown that a 60-s exposure to LWL disrupts brain regions in the dorsal attention network, such as the intraparietal sulcus and the frontal eye fields^92^, and that only SWL – not “orange” light (ʎ_max_ = 590 nm) – modulates the connectivity between a subcortical and cortical brain region involved in regulating attention^93^. Hence, the impact of LWL on alertness and attention remains rather inconclusive in the literature depending on the methodology, including the alertness indicators used, and it remains to be tested whether LWL can activate brain regions in the alerting network to a similar extent as activations associated with exposure to SWL^87,88^.

Melanopic EDI or melanopic irradiance has been shown to predict various NIF responses to light^94^. The SWL condition used in our study was of considerably high m-EDI (∼ 1400 lx), whereas the LWL condition had a low m-EDI (< 5 lx). We therefore hypothesized that SWL would be superior to LWL in improving alertness and attention. However, the results contradicted this hypothesis as there were no differential effects of SWL relative to LWL on any of the PVT measures. Similar findings have been reported by Killgore et al.^34^. One explanation for the lack of observed significant differences between SWL and LWL on PVT outcomes in our study might be that the two narrow bandwidth lights were matched on photon irradiance. When aiming at designing a SWL light condition with a m-EDI (> 1000 lx), matching SWL and LWL on photon irradiance or photon flux results in relatively high L-cone-opic irradiance for LWL. Since melanopsin and other retinal photopigments follow the ‘principle of univariance’, stating that the output of a given photoreceptor is dependent on the total quanta of photons absorbed (i.e., the photoreceptor does not distinguish between changes in wavelength vs. changes in intensity)^19,95^, high-intensity LWL can induce alerting responses as well. Additionally, as retinal opsins show broad spectra tuning, light peaking around e.g., 625 nm (corresponding to “red” LWL) will stimulate not only L-cones but also the other retinal photoreceptors. To be able to conclude mechanistically on the distinct photoreceptor contributions to NIF responses, including alerting effects during the day, metameric light conditions obtained through the method of ‘silent substitution’ are needed^96^. Metameric lights are pairs of light conditions that appear visually similar to an observer but differ in their relative stimulation of the different photoreceptors^97^. Unfortunately, the setup in our laboratory did not enable the administration of metameric lights, and therefore, we cannot draw any mechanistic conclusions in light of our findings. As metameric lights appear visually similar, the use of such light conditions could also have ruled out potential placebo effects arising from participants’ expectations about what light conditions would increase alertness and improve attention. Still, it can be argued that positive and negative effects of light when it comes to health and sleep (e.g., getting enough daylight and avoiding “blue” light exposure in the evening and at night) are more known in public discourse than the neurobehavioral effects of more “extreme” light conditions, such as those of SWL and LWL in our study. Hence, it is unlikely that our results stem from placebo effects alone.

Contrary to one of our main hypotheses, BWL did not induce significant alerting effects despite being of relatively high m-EDI (∼ 1000 lx) and CCT (∼ 8000 K). This finding contrasts results from two other studies testing the effects of broad-spectrum BWL or short-wavelength-enriched white light at a similar time-of-day as that in our study^28,30^. Nevertheless, as shown in a recent systematic review by our lab^57^, most studies report no significant effects of BWL and short-wavelength-enriched white light on PVT measures. It is, however, unclear which parameters of BWL are the most effective in inducing alerting responses during daytime. Although it has been recommended that daytime light should measure at least 250 melanopic lx at eye level^98^, dose-response curves for the acute alerting effects of light during daytime have not yet been firmly established. E.g., Smolders et al.^99^ did not find evidence of any clear dose-response relationships between 1h of exposure to white light (intensity range: ∼ 14-1000 melanopic lx) during daytime on alertness. In addition to the relative proportion of wavelengths in a given light spectrum, dose-response curves for alerting effects of light will also depend on light exposure duration, and the specific time-of-day of exposure, e.g., morning versus afternoon. In the afternoon, following lunch, some experience a dip in core body temperature (i.e., the “post-lunch dip”), which can have negative impacts on performance on cognitive tasks^100^. It is possible that we might have observed alerting effects of BWL with an exposure duration extending into the post-lunch dip^54,101^. Furthermore, the duration of light adaptation was 30 minutes in our study. This duration is similar to other studies in the field, and it aligns with neurophysiological responses to light, showing that the half-maximum EEG gamma response (associated with cognitive processes and attention) is achieved after approx. 12 minutes of light exposure^33^. Still, our results point towards the possibility that a light adaptation longer than 30 minutes prior to cognitive testing might be needed for broad-spectrum BWL to induce improvements in alertness and attention during the day.

Irrespective of light manipulations, prolonged time in testing or repeated test administrations might result in so-called time-on-task effects. On the PVT, this manifests as increasingly slower RTs across administrations^102^. We found evidence of a time-on-task effect with large effect sizes on all PVT metrics (except for RTV), reflected in increasingly slower RTs and an increased number of lapses from measurement block 1 to 2, and from block 1 to 3. In between PVT blocks, participants performed a series of other cognitive and emotion processing tasks, likely contributing to increased fatigue and boredom across time. Initial interaction effect analyses showed that light did not counteract time-on-task effects, a finding that is consistent with results from previous studies^28,48,51^, although significant interactions have been reported as well^12,47^.

To our knowledge, the current study is the first to test whether light influences short-term intra-individual variability in RTs (RTV) as indexed by the SD in RTs across PVT trials^63^. RTV reflects fluctuations in attention and has been proposed as a neurophysiological marker of lapses^103^. Therefore, we expected similar acute alerting effects of light on RTV as on lapses. However, results contradicted this expectation, as there were no significant main effects of light on RTV. Instead, a large proportion of the variance in RTV could be explained by mean RT in our model, with increasingly slower mean RT associated with increasingly higher variability (Pearson’s partial *r* = .66). Additionally, unlike other PVT metrics, RTV did not deteriorate over time. Rather, variability decreased across assessments, indicative of a stable response pattern, likely related to ceiling effects in our young and healthy high-performing sample (see e.g., MacDonald et al.^104^ for a review of changes in RTV across the life span and in relation to brain disorders). Importantly, though, our estimation of RTV might not have been sensitive enough to truly reflect the variability, i.e., the “skewed” (exponential) component of a RT distribution. To model this skewness in particular, ex-gaussian modeling is preferred^63,103,105^. Unfortunately, we did not have enough PVT trials to reliably estimate the exponential component of the RT distribution.

The results of the current study may have been confounded by effects of light on circadian rhythms and sleep. Photic entrainment (phase resetting of the “master” circadian clock, the suprachiasmatic nucleus (SCN)) is achieved by transmittance of light-dark information in the environment through the retinohypothalamic tract originating from melanopsin-containing ipRGCs projecting to the SCN in the hypothalamus^106–108^. The SCN further projects to nuclei involved in the regulation of sleep and alertness/arousal via a pathway connecting the SCN, the dorsomedial hypothalamic nucleus, and the LC^109^. Depending on the timing of light exposure, light can induce either phase-advances if exposure occurs in the morning or phase-delays if exposure occurs in the evening or at night^110–113^. Here, it has been suggested that melanopic illuminance-/irradiance is the best predictor of light’s effects on circadian phase shifting^94^. The presentation order of light conditions in our study was randomized and counter-balanced within groups of participants to reduce confounding related to stronger circadian entrainment and subsequent effects on sleep and alertness following exposure to light with high m-EDI (SWL and BWL). However, we did not impose any control over participants’ light exposure in the evenings. Exposure to artificial light emitted from short-wavelength-enriched LED displays (e.g., from smartphones, tablets, and laptops) or other LED sources in the evening might have induced phase delays and/or exerted negative effects on sleep^114^.

Participants in our study, consisting mainly of university students, underwent cognitive testing at a time of day when alertness levels are typically stable in well-rested individuals. The “intrinsic” (tonic) circadian rhythm of alertness manifests as reduced alertness immediately upon awakening, followed by increasing and rather stable levels throughout the waking day which promotes adequate performance on cognitive tasks^115^. In the evening and at night, circadian rhythms promote decreased alertness and facilitate sleep and rest by interacting with built-up sleep pressure from accumulated wakefulness (i.e., homeostatic processes)^24^. By promoting sleep and rest, circadian and homeostatic factors contribute to improved alertness the day following a consolidated sleep episode^116^. At a group level, studies indicate that university and college students report sleep difficulties and irregular bed and wake times resulting in suboptimal sleep duration when societal demands require relatively early wakeup times^117–120^. When sleep is restricted to a duration below what is optimal for the individual, daytime cognitive functioning is impaired^121^. Even a single night of sleep restriction has been associated with longer RTs and an increased number of lapses of attention^122^. Though we instructed our participants to maintain regular bed and wake times, we did not impose any requirements related to spending a certain amount of time in bed each night. As participants’ sleep duration averaged over the four nights preceding each test session was 6.2±1.2 hours, some of the participants were likely mildly sleep-deprived upon arrival at the laboratory in the morning. Studies indicate that light exposure in the morning following 3-5 hours of nighttime sleep does not improve alertness and performance on challenging cognitive tasks^123,124^, however, sleep deprivation can increase the severity of sleep inertia in the morning^125^. Sleep inertia is a state of reduced arousal and cognitive functioning immediately upon awakening from sleep, which can reduce alertness over the course of a few minutes for up to as long as 3 hours, even in individuals who are not sleep deprived and went to bed at their habitual time^125^. Participants in our study had been awake for an average of 78±32 minutes before entering the laboratory in the morning. Hence, negative cognitive effects of sleep inertia could still have been present for some participants, potentially contributing to enhanced alerting responses to light. Moreover, the effects of morning sleep inertia on cognitive performance depend on chronotype, in that evening chronotypes (with a later circadian timing^126^) display a longer duration of sleep inertia in performance than chronotypes with an earlier circadian timing^127^. Approx. 1 out of 4 participants (25.6%) in our sample self-reported a circadian preference corresponding to a ‘moderately evening’ chronotype. Even though we did not test examine cognitive performance immediately upon awakening, we looked at potential main effects of chronotype on PVT measures as well as interaction effects between chronotype and light on performance but found no significant effects. However, the relatively low sample size in our study likely limited the detection of true null findings. Similarly, limited statistical power likely also reduced our chances of detecting true null effects in the interactions between light condition and season. We did, however, find that participants tested in the fall had fewer lapses than participants tested in spring, and that participants tested in winter had faster optimal responses (10% fastest RTs) than participants tested in summer. These findings suggest that light sensitivity, at least to alerting responses, can have been affected by the length of the photoperiod^43^. However, it is worth pointing out that the number of participants tested in each season was rather small and unbalanced in groups, which could have led to biased estimates in the analyses. As participants were tested in one season only, future research should be conducted with large enough sample sizes enabling the manipulation of season between-subjects or with enough resources enabling repeated testing of the same individual across all four seasons of the year, particularly in studies conducted at high latitudes where there are large seasonal variations in photoperiodic length^65,128,129^.

## Conclusion

The results of the current study show that during exposure to both SWL and LWL in the morning, young, healthy adults were more alert, vigilant, and efficient in their responding, as indicated by faster mean RTs, fewer lapses in attention, and faster optimal responses. Notably, light of substantially high m-EDI was not superior to light of low m-EDI in inducing alerting effects. This finding challenges the assumption that melanopic light quantities are the strongest predictors of a range of NIF responses and indicates that light spectra with high proportions of long wavelengths can also be used to enhance alertness, vigilance, and optimal responding during daytime. Surprisingly, though, despite the fact that the BWL condition had a relatively high m-EDI and high CCT, this light condition did not affect alertness and attention measures, and there were no effects of light on short-term intra-individual RTV. However, future research with higher sample sizes should test whether intra-individual RTV estimated with more optimal techniques such as ex-Gaussian modeling, is acutely affected by exposure to light. Last, although our main findings might have been confounded by the negative effects of suboptimal sleep duration, a study sample consisting of a proportion of acutely, mildly sleep-deprived university students is likely more representative of a “typical” larger population of college/university students, which increases ecological validity and generalizability.

## Data availability

Aggregated behavioral data is available from the corresponding author upon request.

## Author contributions

Writing original draft, planning and conducting statistical analyses, preparing figures and tables: LBB. Methodology and conceptualization: EF-G, EV, IHN, LBB, LS. Funding acquisition: EF-G, IHN, LS. Recruitment of participants: LBB, OBK, MEH, EF-G, EV, IHN. Data collection: LBB, OBK, MEH. Reviewing draft and approving final manuscript for submission: all authors.

## Acknowledgements

We would like to thank Erlend Sunde for assisting with a pre-processing script of PVT raw data, and Mattan S. Ben-Shachar for consulting on multi-level modeling.

