## Supplementary material for "Light of both high and low melanopic illuminance improves alertness and attention during daytime"

#### Supplementary materials

Louise Bruland Bjerrum[1]*, Endre Visted[1], Inger Hilde Nordhus[1], Berge Osnes[1], Bjørn Bjorvatn[2,3], Oda Bugge Kambestad[1], Malika Elise Hansen[1], Lin Sørensen[4] & Elisabeth Flo-Groeneboom[1]

[1]Department of Clinical Psychology, University of Bergen, Norway

[2]Department of Global Public Health and Primary Care, University of Bergen, Norway

[3]Norwegian Competence Center for Sleep Disorders, Haukeland University Hospital, Norway

[4]Department of Biological and Medical Psychology, University of Bergen, Norway

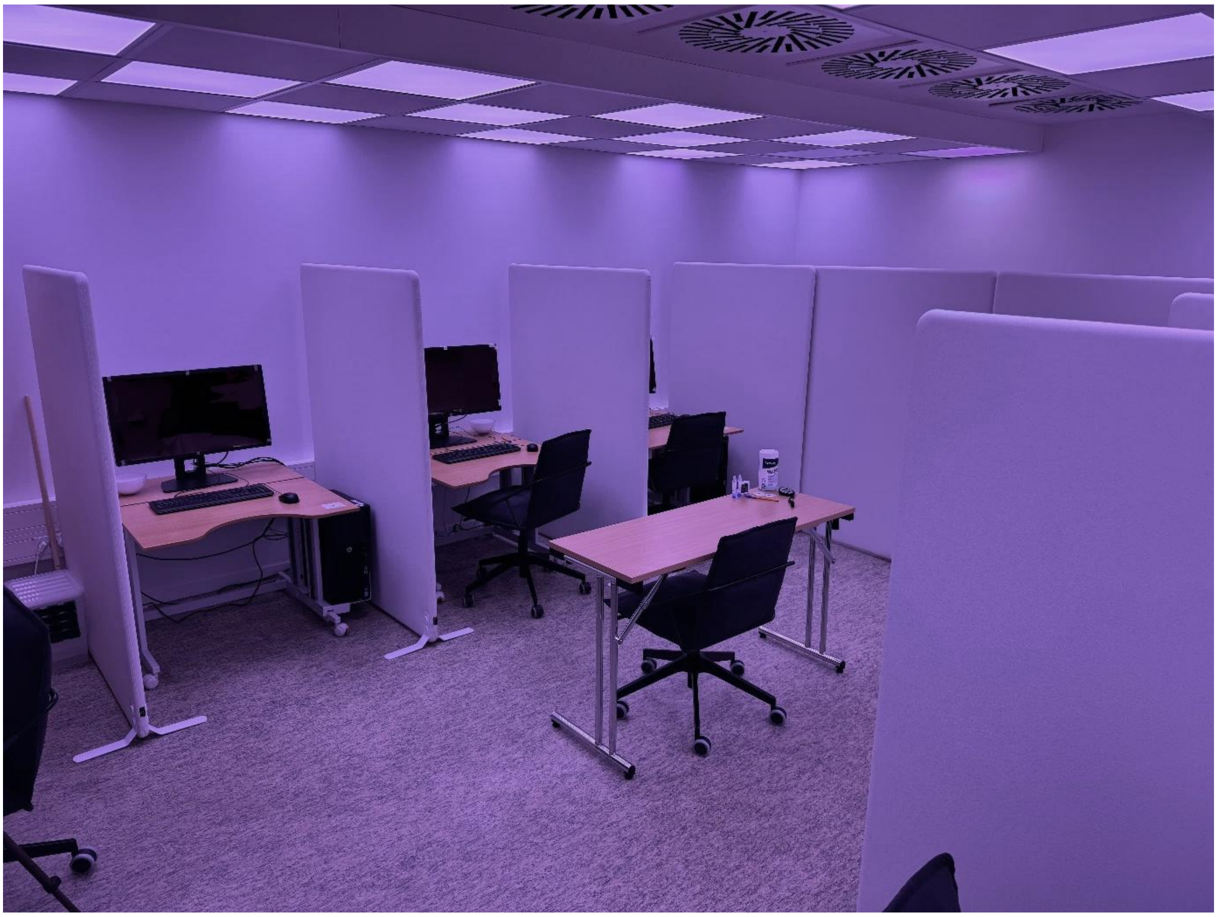

**Figure S1.** Overview of the laboratory setup. The desk in the middle, not separated by partition walls, was used by the researcher carrying out the experiment.

### Light quantities

| | DL<br>(washout) | Narrow bandwidth SWL<br>($\lambda_{max}$ = 455 nm) | Narrow bandwidth LWL<br>($\lambda_{max}$ = 625 nm) | BWL | DL |
|---|---|---|---|---|---|
| **Photopic illuminance, lx** | 1.9<br>(1.7) | 127.8<br>(3.2) | 561.1<br>(13.3) | 983.7<br>(14.3) | 2.2<br>(1.4) |
| **S-cone-opic irradiance, $mW/m^2$** | 4.8<br>(5.8) | 2207.6<br>(53.2) | 2.3<br>(3.9) | 1317.6<br>(20.7) | 5.6<br>(1.8) |
| **M-cone-opic irradiance, $mW/m^2$** | 3.3<br>(4.6) | 565.1<br>(13.9) | 284.6<br>(6.5) | 1353<br>(22) | 6.5<br>(4.2) |
| **L-cone-opic irradiance, $mW/m^2$** | 3.4<br>(2.5) | 323.2<br>(8.0) | 1093.5<br>(26.0) | 1685.7<br>(23.4) | 4.2<br>(2.7) |
| **Rhodopic irradiance, $mW/m^2$** | 5.7<br>(7.5) | 1554<br>(38) | 27.3<br>(1.6) | 1526<br>(23.9) | 12.1<br>(7.9) |
| **Melanopic irradiance, $mW/m^2$** | 6.3<br>(8.2) | 1912.1<br>(46.7) | 5.0<br>(1.7) | 1532.8<br>(23.3) | 12.9<br>(8.5) |
| **S-cone-opic EDI, lx** | 5.9<br>(7.2) | 2701.1<br>(65.2) | 2.9<br>(4.7) | 1612.1<br>(25.3) | 6.9<br>(2.2) |
| **M-cone-opic EDI, lx** | 2.3<br>(3.2) | 388.2<br>(9.6) | 195.5<br>(4.5) | 929.3<br>(15.1) | 4.5<br>(2.9) |
| **L-cone-opic EDI, lx** | 2.1<br>(1.5) | 198.4<br>(4.9) | 671.3<br>(16) | 1034.9<br>(14.4) | 2.6<br>(1.7) |
| **Rhodopic EDI, lx** | 3.9<br>(5.2) | 1072<br>(26.2) | 18.9<br>(1.1) | 1052.6<br>(16.5) | 8.4<br>(5.4) |
| **Melanopic EDI, lx** | 4.8<br>(6.2) | 1441.8<br>(35.2) | 3.8<br>(1.3) | 1155.8<br>(17.6) | 9.7<br>(6.4) |
| **S-cone-opic ELR** | 1.9 | 17.3 | 0 | 1.3 | 39.1 |
| **M-cone-opic ELR** | 1.1 | 4.4 | 0.5 | 1.4 | 4.5 |
| **L-cone-opic ELR** | 2 | 2.5 | 1.9 | 1.7 | 3.4 |
| **Rhodopic ELR** | 2 | 12.2 | 0 | 1.6 | 12.3 |
| **Melanopic ELR** | 2.3 | 15 | 0 | 1.6 | 15.3 |
| **CCT, K** | | | | 8035.7<br>(42.8) | |

**Table S1.** α-opic light quantities at eye level for each light condition. Light stimuli were measured at eye level (vertically, 120 cm above the floor, ~90 cm horizontally from the computer screen in the direction of gaze). All light quantities were calculated using luox[1,2]. Values are in mean (sd). BWL = Bright White Light. CCT = Correlated Color Temperature. DL = Dim Light. 'DL washout' refers to the 1-hour exposure to DL preceding light adaptation. EDI = Equivalent Daylight Illuminance. ELR = Efficiency of Luminous Radiation. K = Kelvin. LWL = Long-Wavelength Light. lx = lux. nm = nanometer. SWL = Short-Wavelength Light.

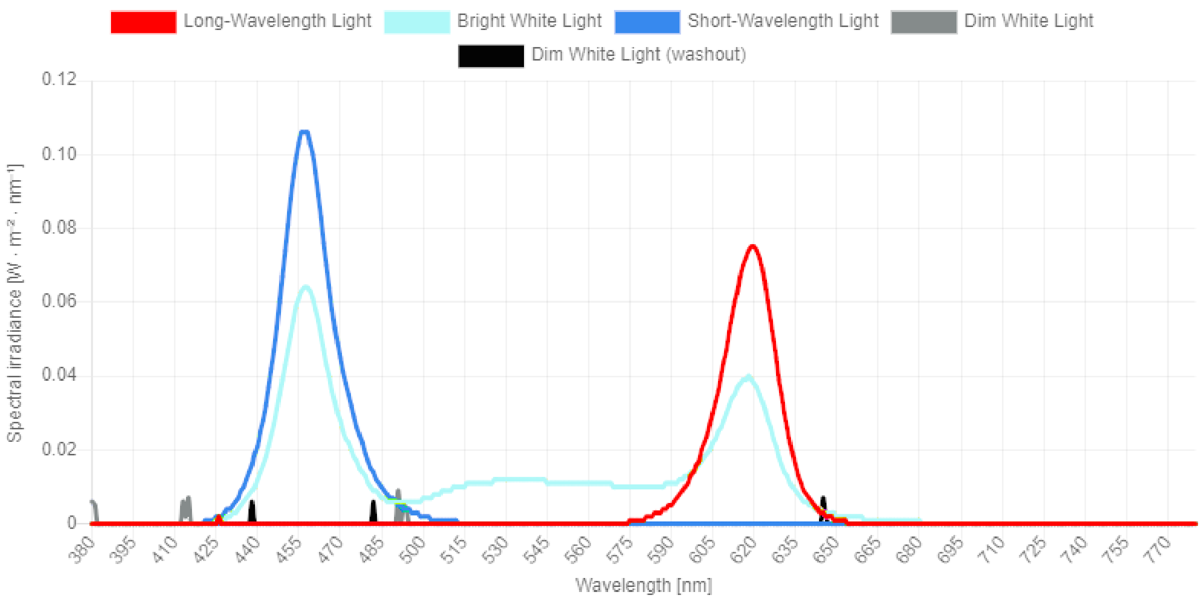


**Figure S2.** Spectral power distribution curves in each light condition, obtained from *luox*[1,2]. 'Dim White Light (washout)' refers to the 1-hour of exposure to dim light preceding light adaptation. nm = nanometer.

| Wavelength (nm) | LWL | BWL | SWL | DL | DL (washout) |
|---|---|---|---|---|---|
| 380 | 0,0131242 | 0,0123232 | 0,00167491 | 6,2867e05 | 0,0108774 |
| 381 | 0,0239545 | 0,0029097 | 0,00160094 | 5,65926e06 | 0,0100918 |
| 382 | 0,0370853 | 0,0201698 | 0,00849849 | 0,000138825 | 0,00912829 |
| 383 | 0,0436829 | 0,0230918 | 0,00792982 | 0,00133805 | 0,0080723 |
| 384 | 0,045514 | 0,015553 | 0,0413762 | 0,0033068 | 0,00694886 |
| 385 | 0,0451451 | 0,00804538 | 0,0621028 | 0,00476319 | 0,00609343 |
| 386 | 0,0375037 | 0,000640634 | 0,0407813 | 0,00452587 | 0,00612393 |
| 387 | 0,0298623 | 0,00676411 | 0,0194599 | 0,00428854 | 0,00615443 |
| 388 | 0,0392953 | 0,00683554 | 0,0150067 | 0,00323292 | 0,00708467 |
| 389 | 0,0508677 | 0,023067 | 0,0126671 | 0,00207477 | 0,00812765 |
| 390 | 0,055362 | 0,0194049 | 0,0047153 | 0,00146008 | 0,008665 |
| 391 | 0,0540049 | 0,000703127 | 0,0078762 | 0,00129466 | 0,00878435 |
| 392 | 0,0517059 | 0,0133018 | 0,0140263 | 0,00100027 | 0,00897256 |
| 393 | 0,0458342 | 0,00258303 | 0,00425655 | 0,000216729 | 0,00942197 |
| 394 | 0,0399626 | 0,0184679 | 0,0225394 | 0,000566816 | 0,00987139 |
| 395 | 0,0525091 | 0,00999464 | 0,040846 | 0,00109264 | 0,00799911 |
| 396 | 0,0677521 | 0,00204472 | 0,059156 | 0,00310975 | 0,00578692 |
| 397 | 0,0696787 | 0,00881675 | 0,0710473 | 0,00339913 | 0,00563297 |
| 398 | 0,0601909 | 0,0110738 | 0,0774367 | 0,00220756 | 0,00724326 |
| 399 | 0,0511872 | 0,0128833 | 0,0824601 | 0,00118677 | 0,00857948 |
| 400 | 0,0440359 | 0,0129799 | 0,0822556 | 0,000819547 | 0,00886686 |
| 401 | 0,0368846 | 0,0130764 | 0,0820511 | 0,000452325 | 0,00915423 |
| 402 | 0,0348866 | 0,0161463 | 0,0712498 | 0,000568316 | 0,00881925 |
| 403 | 0,0336015 | 0,0196276 | 0,0589823 | 0,000751161 | 0,00839817 |
| 404 | 0,0331578 | 0,027506 | 0,0567647 | 0,000794978 | 0,00807166 |
| 405 | 0,0333959 | 0,0389475 | 0,0626908 | 0,000726139 | 0,00782179 |
| 406 | 0,0342842 | 0,0506982 | 0,0690533 | 0,000861671 | 0,00748311 |
| 407 | 0,0373618 | 0,0634902 | 0,0768856 | 0,00168535 | 0,00684537 |
| 408 | 0,0404394 | 0,0762823 | 0,0847178 | 0,00250904 | 0,00620763 |
| 409 | 0,0250981 | 0,0796647 | 0,10403 | 0,00221209 | 0,00645532 |
| 410 | 0,00789678 | 0,082097 | 0,124502 | 0,00180198 | 0,00679242 |
| 411 | 0,00506341 | 0,094687 | 0,156041 | 0,00123653 | 0,00742349 |
| 412 | 0,0122009 | 0,114326 | 0,195261 | 0,000563285 | 0,00825855 |
| 413 | 0,0180907 | 0,135039 | 0,240505 | 6,52774e05 | 0,00885405 |
| 414 | 0,0206943 | 0,158579 | 0,301619 | 2,88867e05 | 0,00881852 |
| 415 | 0,0232979 | 0,18212 | 0,362734 | 7,50402e06 | 0,00878298 |
| 416 | 0,0230382 | 0,22403 | 0,438839 | 0,00118236 | 0,00747115 |
| 417 | 0,0226679 | 0,266649 | 0,515524 | 0,00241957 | 0,00611005 |
| 418 | 0,0173217 | 0,329627 | 0,603674 | 0,00274627 | 0,00539151 |
| 419 | 0,00933541 | 0,403407 | 0,697908 | 0,00258989 | 0,00501389 |
| 420 | 0,00827722 | 0,487073 | 0,821916 | 0,00230732 | 0,00515498 |
| 421 | 0,0202693 | 0,589361 | 1,00201 | 0,00178705 | 0,00627315 |
| 422 | 0,0313725 | 0,692882 | 1,18458 | 0,00124945 | 0,00734762 |
| 423 | 0,0227601 | 0,823775 | 1,42208 | 0,000327524 | 0,00745292 |

| | | | | | |
|---|---|---|---|---|---|
| 424 | 0,0141478 | 0,954667 | 1,65958 | 0,000594398 | 0,00755823 |
| 425 | 0,00685745 | 1,15305 | 2,01144 | 0,000176471 | 0,0067088 |
| 426 | 2,56991e05 | 1,37483 | 2,40296 | 0,000706169 | 0,00552823 |
| 427 | 0,00115814 | 1,63646 | 2,87185 | 0,00128118 | 0,00483099 |
| 428 | 0,0122304 | 1,94781 | 3,43729 | 0,00147224 | 0,00473698 |
| 429 | 0,021144 | 2,2796 | 4,03694 | 0,00155288 | 0,00470657 |
| 430 | 0,0178472 | 2,72701 | 4,83012 | 0,00100888 | 0,00503597 |
| 431 | 0,0145504 | 3,17442 | 5,6233 | 0,000464886 | 0,00536536 |
| 432 | 0,0103596 | 3,78465 | 6,68093 | 0,00063378 | 0,00478038 |
| 433 | 0,00602174 | 4,42168 | 7,78207 | 0,000919981 | 0,00404494 |
| 434 | 0,00672155 | 5,19537 | 9,12257 | 0,00102865 | 0,00403164 |
| 435 | 0,0112172 | 6,07204 | 10,6434 | 0,00100356 | 0,00456247 |
| 436 | 0,0166848 | 7,03351 | 12,3056 | 0,00130897 | 0,0043337 |
| 437 | 0,0245907 | 8,2077 | 14,3224 | 0,00244338 | 0,00219966 |
| 438 | 0,0324433 | 9,38308 | 16,3413 | 0,00357528 | 6,66142e05 |
| 439 | 0,0223972 | 10,9579 | 19,0752 | 0,00386585 | 0,00173457 |
| 440 | 0,0123512 | 12,5327 | 21,8091 | 0,00415643 | 0,00353576 |
| 441 | 0,012711 | 14,4457 | 25,1189 | 0,00326424 | 0,00333256 |
| 442 | 0,0170354 | 16,4876 | 28,648 | 0,00192145 | 0,00236572 |
| 443 | 0,0247094 | 18,8161 | 32,6664 | 0,00102396 | 0,00196618 |
| 444 | 0,0365019 | 21,497 | 37,2864 | 0,000674001 | 0,00226416 |
| 445 | 0,0466736 | 24,3049 | 42,1161 | 0,000492281 | 0,00282673 |
| 446 | 0,0492676 | 27,7066 | 47,926 | 0,0010971 | 0,00462626 |
| 447 | 0,0518616 | 31,1084 | 53,736 | 0,00170191 | 0,0064258 |
| 448 | 0,0542345 | 35,1115 | 60,473 | 0,00129752 | 0,00668565 |
| 449 | 0,0565848 | 39,176 | 67,3049 | 0,000789905 | 0,00678803 |
| 450 | 0,0568628 | 43,4115 | 74,3237 | 0,00107141 | 0,00768482 |
| 451 | 0,0559831 | 47,7425 | 81,447 | 0,00179379 | 0,00902545 |
| 452 | 0,0536605 | 51,871 | 88,1536 | 0,00226041 | 0,00988798 |
| 453 | 0,0489767 | 55,6681 | 94,1784 | 0,0023085 | 0,0099681 |
| 454 | 0,044099 | 59,2803 | 99,8761 | 0,00227896 | 0,00995986 |
| 455 | 0,0378051 | 61,5428 | 103,184 | 0,0016825 | 0,00930618 |
| 456 | 0,0315112 | 63,8053 | 106,492 | 0,00108605 | 0,00865251 |
| 457 | 0,0385491 | 64,2267 | 106,657 | 0,00150659 | 0,00888961 |
| 458 | 0,0476672 | 64,3607 | 106,331 | 0,00208584 | 0,00926572 |
| 459 | 0,0475729 | 63,1578 | 103,848 | 0,00252636 | 0,00957814 |
| 460 | 0,0416642 | 61,111 | 100,004 | 0,00287932 | 0,00985035 |
| 461 | 0,0365078 | 58,5355 | 95,3558 | 0,00302282 | 0,00987801 |
| 462 | 0,0326623 | 55,0387 | 89,3065 | 0,00280133 | 0,00947954 |
| 463 | 0,0305101 | 51,4885 | 83,2004 | 0,0026098 | 0,00908112 |
| 464 | 0,0408093 | 47,5457 | 76,677 | 0,0026386 | 0,00868311 |
| 465 | 0,0511085 | 43,6029 | 70,1535 | 0,00266741 | 0,0082851 |
| 466 | 0,0558025 | 40,0334 | 64,335 | 0,00216113 | 0,00670841 |
| 467 | 0,0597203 | 36,5155 | 58,6141 | 0,00158075 | 0,00496849 |
| 468 | 0,056942 | 33,462 | 53,674 | 0,0018533 | 0,0046408 |

| | | | | | |
|---|---|---|---|---|---|
| 469 | 0,050407 | 30,669 | 49,1719 | 0,0026044 | 0,00510545 |
| 470 | 0,0445984 | 28,1379 | 45,0662 | 0,00315638 | 0,0054907 |
| 471 | 0,0398507 | 25,9894 | 41,5395 | 0,0034176 | 0,00575999 |
| 472 | 0,0354208 | 23,9023 | 38,112 | 0,00348288 | 0,00559746 |
| 473 | 0,0324796 | 22,1028 | 35,1497 | 0,00263047 | 0,00341237 |
| 474 | 0,0295385 | 20,3033 | 32,1873 | 0,00177806 | 0,00122728 |
| 475 | 0,0311041 | 18,7165 | 29,5179 | 0,00177917 | 0,00109017 |
| 476 | 0,0329114 | 17,141 | 26,8641 | 0,00182603 | 0,00106288 |
| 477 | 0,0353187 | 15,7065 | 24,4212 | 0,00201272 | 0,000870729 |
| 478 | 0,0379525 | 14,3252 | 22,0579 | 0,00225219 | 0,00061634 |
| 479 | 0,0371366 | 13,0528 | 19,8623 | 0,00221133 | 0,000265568 |
| 480 | 0,0329603 | 11,8866 | 17,8299 | 0,0018974 | 0,000179092 |
| 481 | 0,0301532 | 10,8003 | 15,9235 | 0,00175308 | 0,000381489 |
| 482 | 0,0306839 | 9,90861 | 14,3242 | 0,0020222 | 6,65519e06 |
| 483 | 0,0314935 | 9,03819 | 12,7563 | 0,00228638 | 0,000297374 |
| 484 | 0,0355535 | 8,41534 | 11,5543 | 0,00249288 | 0,000547438 |
| 485 | 0,0396135 | 7,79249 | 10,3522 | 0,00269939 | 0,00139225 |
| 486 | 0,0457386 | 7,36679 | 9,40704 | 0,00192366 | 0,00180155 |
| 487 | 0,0521697 | 6,97029 | 8,49997 | 0,00100238 | 0,0021463 |
| 488 | 0,0458673 | 6,69797 | 7,75424 | 0,00068976 | 0,00248009 |
| 489 | 0,0331985 | 6,48773 | 7,08917 | 0,000681442 | 0,00280838 |
| 490 | 0,0262791 | 6,33447 | 6,48108 | 0,000491416 | 0,00329313 |
| 491 | 0,0259406 | 6,24644 | 5,93821 | 9,34041e05 | 0,00395693 |
| 492 | 0,0278406 | 6,1833 | 5,41583 | 0,000168022 | 0,00427058 |
| 493 | 0,0358002 | 6,18751 | 4,94892 | 5,97107e05 | 0,00363632 |
| 494 | 0,0419916 | 6,19831 | 4,48881 | 4,51094e05 | 0,0030317 |
| 495 | 0,027027 | 6,28789 | 4,11 | 0,000108157 | 0,00278159 |
| 496 | 0,0120625 | 6,37748 | 3,73119 | 0,000171205 | 0,00253148 |
| 497 | 0,016783 | 6,53412 | 3,39983 | 0,000700481 | 0,00266439 |
| 498 | 0,0239753 | 6,69918 | 3,07442 | 0,0012883 | 0,00284541 |
| 499 | 0,0232206 | 6,90232 | 2,80821 | 0,00110674 | 0,00317932 |
| 500 | 0,0191196 | 7,12149 | 2,56694 | 0,000601188 | 0,00357761 |
| 501 | 0,0206217 | 7,36688 | 2,34542 | 0,000183416 | 0,00422616 |
| 502 | 0,0272361 | 7,63619 | 2,14194 | 0,000154268 | 0,00510305 |
| 503 | 0,0290528 | 7,91282 | 1,95965 | 0,000160725 | 0,00523968 |
| 504 | 0,0218055 | 8,20328 | 1,8174 | 0,000458582 | 0,00397781 |
| 505 | 0,0167088 | 8,49256 | 1,67815 | 0,0010205 | 0,00289251 |
| 506 | 0,0218682 | 8,77623 | 1,55315 | 0,0013087 | 0,00264927 |
| 507 | 0,0270011 | 9,05982 | 1,42834 | 0,00159655 | 0,00240337 |
| 508 | 0,0264012 | 9,32806 | 1,34413 | 0,00180763 | 0,00158164 |
| 509 | 0,0258013 | 9,5963 | 1,25991 | 0,0020187 | 0,000759909 |
| 510 | 0,0229949 | 9,84506 | 1,18043 | 0,00174424 | 0,00114211 |
| 511 | 0,0197681 | 10,0901 | 1,10185 | 0,00137725 | 0,00175377 |
| 512 | 0,0182672 | 10,3179 | 1,0347 | 0,00163974 | 0,00143117 |
| 513 | 0,0175811 | 10,5376 | 0,972952 | 0,00219937 | 0,000667559 |

| | | | | | |
|---|---|---|---|---|---|
| 514 | 0,0206023 | 10,7336 | 0,919235 | 0,00261919 | 0,000558887 |
| 515 | 0,0270072 | 10,9079 | 0,872849 | 0,0029114 | 0,00104797 |
| 516 | 0,0311103 | 11,075 | 0,824886 | 0,00326583 | 0,00122765 |
| 517 | 0,0313014 | 11,23 | 0,774244 | 0,00372601 | 0,000881532 |
| 518 | 0,0318817 | 11,3759 | 0,727968 | 0,00419624 | 0,000612628 |
| 519 | 0,0338236 | 11,49 | 0,696976 | 0,00470166 | 0,000613967 |
| 520 | 0,0358305 | 11,6023 | 0,666201 | 0,00530053 | 0,000557879 |
| 521 | 0,0386031 | 11,6943 | 0,637987 | 0,007002 | 0,000175697 |
| 522 | 0,0413757 | 11,7863 | 0,609773 | 0,00870346 | 0,000909273 |
| 523 | 0,0380988 | 11,8517 | 0,598098 | 0,0101182 | 0,000819743 |
| 524 | 0,0344237 | 11,9153 | 0,587511 | 0,011514 | 0,000676039 |
| 525 | 0,0430176 | 11,9615 | 0,563819 | 0,0136609 | 0,000626427 |
| 526 | 0,05463 | 12,0034 | 0,536903 | 0,0159925 | 0,000599963 |
| 527 | 0,0600338 | 12,0398 | 0,518238 | 0,0184593 | 0,000914789 |
| 528 | 0,0623947 | 12,0735 | 0,503617 | 0,0209924 | 0,00139689 |
| 529 | 0,0690005 | 12,0978 | 0,499412 | 0,0244488 | 0,00235588 |
| 530 | 0,0791669 | 12,114 | 0,503942 | 0,0286797 | 0,00371485 |
| 531 | 0,0830038 | 12,1258 | 0,499061 | 0,0325691 | 0,00452693 |
| 532 | 0,0781311 | 12,1315 | 0,481229 | 0,0359886 | 0,00458647 |
| 533 | 0,0767019 | 12,1342 | 0,470369 | 0,0395059 | 0,00466758 |
| 534 | 0,0832253 | 12,1299 | 0,475611 | 0,0432491 | 0,0047985 |
| 535 | 0,0878296 | 12,1264 | 0,475231 | 0,0470708 | 0,00491813 |
| 536 | 0,0841313 | 12,1263 | 0,450529 | 0,0512315 | 0,0049889 |
| 537 | 0,0817703 | 12,1255 | 0,428103 | 0,0554262 | 0,00506879 |
| 538 | 0,0952952 | 12,1164 | 0,432725 | 0,0600235 | 0,00525701 |
| 539 | 0,10882 | 12,1073 | 0,437347 | 0,0646209 | 0,00544523 |
| 540 | 0,120046 | 12,0988 | 0,438612 | 0,0680046 | 0,00495413 |
| 541 | 0,131205 | 12,0902 | 0,439781 | 0,0713532 | 0,00444336 |
| 542 | 0,128293 | 12,0701 | 0,434535 | 0,0760183 | 0,00550426 |
| 543 | 0,123286 | 12,0482 | 0,428334 | 0,0808793 | 0,00679911 |
| 544 | 0,130269 | 12,0214 | 0,431569 | 0,084304 | 0,00509146 |
| 545 | 0,140766 | 11,9931 | 0,43757 | 0,0873079 | 0,00250387 |
| 546 | 0,147155 | 11,9675 | 0,437475 | 0,0909774 | 0,00236548 |
| 547 | 0,151616 | 11,9432 | 0,434518 | 0,0949594 | 0,00337622 |
| 548 | 0,157494 | 11,918 | 0,427266 | 0,0985348 | 0,00387965 |
| 549 | 0,164348 | 11,8922 | 0,417058 | 0,101831 | 0,00403395 |
| 550 | 0,168523 | 11,8615 | 0,409735 | 0,105607 | 0,0050083 |
| 551 | 0,170108 | 11,826 | 0,4052 | 0,109848 | 0,00677547 |
| 552 | 0,173482 | 11,7899 | 0,395836 | 0,113128 | 0,00735492 |
| 553 | 0,179237 | 11,753 | 0,380043 | 0,115129 | 0,00635332 |
| 554 | 0,184119 | 11,7145 | 0,376897 | 0,117111 | 0,00553724 |
| 555 | 0,187406 | 11,673 | 0,396834 | 0,119061 | 0,00505981 |
| 556 | 0,191186 | 11,6302 | 0,409257 | 0,121064 | 0,00500121 |
| 557 | 0,19621 | 11,5842 | 0,402675 | 0,123201 | 0,00600197 |
| 558 | 0,204257 | 11,5382 | 0,399752 | 0,124927 | 0,00655591 |

| | | | | | |
|---|---|---|---|---|---|
| 559 | 0,223199 | 11,4928 | 0,410013 | 0,12517 | 0,00549975 |
| 560 | 0,240529 | 11,4455 | 0,417519 | 0,125546 | 0,00469469 |
| 561 | 0,249113 | 11,3887 | 0,410078 | 0,126643 | 0,00525239 |
| 562 | 0,259824 | 11,3322 | 0,404164 | 0,127674 | 0,00570407 |
| 563 | 0,290017 | 11,2783 | 0,412237 | 0,128099 | 0,00518487 |
| 564 | 0,320389 | 11,2241 | 0,419122 | 0,128515 | 0,00471786 |
| 565 | 0,354486 | 11,1663 | 0,401267 | 0,128756 | 0,0053377 |
| 566 | 0,388584 | 11,1084 | 0,383412 | 0,128996 | 0,00595755 |
| 567 | 0,44037 | 11,0572 | 0,402939 | 0,128751 | 0,00521015 |
| 568 | 0,492196 | 11,006 | 0,422552 | 0,128505 | 0,00445962 |
| 569 | 0,55329 | 10,9434 | 0,40192 | 0,128461 | 0,00466981 |
| 570 | 0,61483 | 10,8802 | 0,379354 | 0,128426 | 0,00492616 |
| 571 | 0,687448 | 10,8346 | 0,382964 | 0,127765 | 0,00469098 |
| 572 | 0,761093 | 10,7907 | 0,389001 | 0,127045 | 0,00441023 |
| 573 | 0,86541 | 10,7498 | 0,380719 | 0,126508 | 0,00411406 |
| 574 | 0,973883 | 10,7094 | 0,370498 | 0,125996 | 0,00381581 |
| 575 | 1,1019 | 10,6762 | 0,37074 | 0,125511 | 0,00402949 |
| 576 | 1,23333 | 10,6442 | 0,372818 | 0,12503 | 0,00433295 |
| 577 | 1,39524 | 10,6235 | 0,369805 | 0,124059 | 0,00407258 |
| 578 | 1,56358 | 10,6052 | 0,365716 | 0,122985 | 0,00369304 |
| 579 | 1,75228 | 10,6013 | 0,360431 | 0,121495 | 0,00252936 |
| 580 | 1,94592 | 10,6008 | 0,354856 | 0,119905 | 0,00117551 |
| 581 | 2,17682 | 10,6163 | 0,354675 | 0,118635 | 0,000112837 |
| 582 | 2,41771 | 10,6362 | 0,35594 | 0,117451 | 0,000871846 |
| 583 | 2,72731 | 10,6728 | 0,356633 | 0,117438 | 0,000452928 |
| 584 | 3,0566 | 10,7141 | 0,357162 | 0,117762 | 0,00243944 |
| 585 | 3,42867 | 10,7876 | 0,34824 | 0,11659 | 0,00186562 |
| 586 | 3,81348 | 10,8707 | 0,336503 | 0,114974 | 0,00052904 |
| 587 | 4,28388 | 10,9819 | 0,340171 | 0,114695 | 0,00168712 |
| 588 | 4,78009 | 11,1015 | 0,348486 | 0,114818 | 0,00359758 |
| 589 | 5,35486 | 11,2701 | 0,33733 | 0,11427 | 0,0039535 |
| 590 | 5,95299 | 11,4534 | 0,320385 | 0,113522 | 0,00384711 |
| 591 | 6,67131 | 11,703 | 0,320106 | 0,11293 | 0,00393677 |
| 592 | 7,42393 | 11,9717 | 0,324584 | 0,112384 | 0,0040824 |
| 593 | 8,33363 | 12,3001 | 0,319728 | 0,110773 | 0,0032286 |
| 594 | 9,28514 | 12,6444 | 0,312387 | 0,108879 | 0,00210873 |
| 595 | 10,4248 | 13,0882 | 0,303149 | 0,108133 | 0,00202514 |
| 596 | 11,6098 | 13,5558 | 0,293456 | 0,107662 | 0,00219055 |
| 597 | 13,012 | 14,1292 | 0,282705 | 0,106525 | 0,0016743 |
| 598 | 14,4596 | 14,7247 | 0,271735 | 0,105249 | 0,00101595 |
| 599 | 16,1672 | 15,5018 | 0,260964 | 0,104138 | 0,00108231 |
| 600 | 17,9195 | 16,3101 | 0,250228 | 0,103056 | 0,00127315 |
| 601 | 20,0287 | 17,267 | 0,24964 | 0,101875 | 0,00108752 |
| 602 | 22,1848 | 18,2434 | 0,250384 | 0,100681 | 0,000852527 |
| 603 | 24,7197 | 19,4394 | 0,255836 | 0,099863 | 0,00156846 |

| | | | | | |
|---|---|---|---|---|---|
| 604 | 27,2878 | 20,6547 | 0,2617 | 0,099078 | 0,00236771 |
| 605 | 30,2883 | 22,0756 | 0,247254 | 0,097332 | 0,00187402 |
| 606 | 33,3089 | 23,5061 | 0,231977 | 0,0955511 | 0,00133444 |
| 607 | 36,7637 | 25,1486 | 0,222518 | 0,0950098 | 0,00273326 |
| 608 | 40,24 | 26,7992 | 0,214323 | 0,0944391 | 0,00402663 |
| 609 | 44,0919 | 28,5922 | 0,228246 | 0,0933533 | 0,00347386 |
| 610 | 47,9675 | 30,3884 | 0,239731 | 0,0922447 | 0,00296915 |
| 611 | 52,0384 | 32,2113 | 0,231172 | 0,0909485 | 0,00285971 |
| 612 | 56,0969 | 34,0028 | 0,222316 | 0,0897108 | 0,00280563 |
| 613 | 60,0932 | 35,6384 | 0,211992 | 0,0887633 | 0,00302656 |
| 614 | 63,9666 | 37,1743 | 0,20362 | 0,0881098 | 0,00362157 |
| 615 | 67,4319 | 38,3792 | 0,201726 | 0,0884323 | 0,00545836 |
| 616 | 70,5733 | 39,3715 | 0,200182 | 0,0884102 | 0,00650413 |
| 617 | 72,9597 | 39,8683 | 0,199454 | 0,0875852 | 0,00570627 |
| 618 | 74,7434 | 40,0353 | 0,192267 | 0,0864878 | 0,00432018 |
| 619 | 75,5172 | 39,6497 | 0,174258 | 0,0849338 | 0,00194851 |
| 620 | 75,4306 | 38,8421 | 0,168642 | 0,083967 | 0,000853253 |
| 621 | 74,3013 | 37,523 | 0,178045 | 0,0837119 | 0,00130493 |
| 622 | 72,155 | 35,8053 | 0,17691 | 0,0832246 | 0,00190072 |
| 623 | 69,1256 | 33,7415 | 0,166625 | 0,0825356 | 0,00262166 |
| 624 | 65,2049 | 31,403 | 0,167882 | 0,0813171 | 0,00192772 |
| 625 | 60,7454 | 28,8986 | 0,176119 | 0,0797784 | 0,000378339 |
| 626 | 55,7413 | 26,3267 | 0,181892 | 0,0807354 | 0,00286416 |
| 627 | 50,5212 | 23,7282 | 0,186686 | 0,0826823 | 0,0069505 |
| 628 | 45,2287 | 21,2801 | 0,189495 | 0,0812672 | 0,00660693 |
| 629 | 39,9197 | 18,8665 | 0,191848 | 0,0790821 | 0,0052487 |
| 630 | 35,082 | 16,7919 | 0,182021 | 0,0792093 | 0,00613709 |
| 631 | 30,3051 | 14,7585 | 0,171076 | 0,0795481 | 0,00724018 |
| 632 | 26,2635 | 13,1502 | 0,159976 | 0,0798751 | 0,00871775 |
| 633 | 22,3436 | 11,5891 | 0,149606 | 0,0799922 | 0,00967784 |
| 634 | 19,1998 | 10,329 | 0,143876 | 0,0787721 | 0,00733994 |
| 635 | 16,2431 | 9,14779 | 0,143645 | 0,0777733 | 0,00562179 |
| 636 | 13,8396 | 8,20021 | 0,159675 | 0,077429 | 0,00573662 |
| 637 | 11,6733 | 7,34709 | 0,165858 | 0,0771201 | 0,00558047 |
| 638 | 9,90303 | 6,65167 | 0,155598 | 0,0768704 | 0,00497192 |
| 639 | 8,38242 | 6,04354 | 0,145841 | 0,0765794 | 0,00431939 |
| 640 | 7,11009 | 5,52221 | 0,136582 | 0,0762473 | 0,00362312 |
| 641 | 6,05608 | 5,08508 | 0,140701 | 0,0753928 | 0,00239722 |
| 642 | 5,12861 | 4,69674 | 0,152575 | 0,0742354 | 0,000864352 |
| 643 | 4,37926 | 4,36087 | 0,148911 | 0,0735286 | 0,000221652 |
| 644 | 3,68322 | 4,0407 | 0,140597 | 0,0729565 | 0,00015463 |
| 645 | 3,19333 | 3,80961 | 0,134916 | 0,0727508 | 7,22134e05 |
| 646 | 2,72985 | 3,5887 | 0,129941 | 0,0724867 | 2,70306e05 |
| 647 | 2,36972 | 3,39227 | 0,13233 | 0,0706786 | 0,00132595 |
| 648 | 2,03262 | 3,20553 | 0,130654 | 0,0691599 | 0,00235631 |

| | | | | | |
|---|---|---|---|---|---|
| 649 | 1,7828 | 3,05552 | 0,113573 | 0,0687385 | 0,00236877 |
| 650 | 1,5552 | 2,91578 | 0,0969539 | 0,067823 | 0,00292976 |
| 651 | 1,36625 | 2,79388 | 0,0811404 | 0,066048 | 0,00444525 |
| 652 | 1,19708 | 2,67405 | 0,0800822 | 0,0649511 | 0,00521837 |
| 653 | 1,04573 | 2,55607 | 0,0923174 | 0,0644652 | 0,00532268 |
| 654 | 0,91451 | 2,45521 | 0,10366 | 0,0635845 | 0,00507331 |
| 655 | 0,792251 | 2,36198 | 0,114605 | 0,0625282 | 0,0046666 |
| 656 | 0,718517 | 2,27826 | 0,105029 | 0,0620423 | 0,00440378 |
| 657 | 0,652031 | 2,19663 | 0,0923181 | 0,0616454 | 0,0041817 |
| 658 | 0,574402 | 2,12935 | 0,0826191 | 0,0612114 | 0,00440162 |
| 659 | 0,505145 | 2,06055 | 0,0760232 | 0,0608546 | 0,00410925 |
| 660 | 0,465609 | 1,98634 | 0,0804422 | 0,0607716 | 0,00199832 |
| 661 | 0,425312 | 1,91367 | 0,0826697 | 0,0595968 | 0,00203555 |
| 662 | 0,3839 | 1,84325 | 0,0816852 | 0,0568219 | 0,00522129 |
| 663 | 0,35128 | 1,77949 | 0,0728275 | 0,0553945 | 0,00705328 |
| 664 | 0,324613 | 1,72026 | 0,0586397 | 0,0548794 | 0,00796878 |
| 665 | 0,296937 | 1,66792 | 0,0546594 | 0,0553248 | 0,00578307 |
| 666 | 0,268995 | 1,6174 | 0,0533602 | 0,0560226 | 0,00278283 |
| 667 | 0,241331 | 1,55365 | 0,0519983 | 0,0552339 | 0,00287162 |
| 668 | 0,218687 | 1,49326 | 0,0510106 | 0,0542175 | 0,00316996 |
| 669 | 0,216336 | 1,44688 | 0,0515385 | 0,0523245 | 0,00422306 |
| 670 | 0,214003 | 1,40373 | 0,0444873 | 0,051662 | 0,00402961 |
| 671 | 0,211698 | 1,36524 | 0,0264763 | 0,0527787 | 0,00203359 |
| 672 | 0,190809 | 1,32339 | 0,0396613 | 0,0510703 | 0,00351221 |
| 673 | 0,158741 | 1,27952 | 0,0716141 | 0,0476621 | 0,0070812 |
| 674 | 0,143967 | 1,22916 | 0,0444107 | 0,0474077 | 0,00563576 |
| 675 | 0,133677 | 1,1798 | 0,00966137 | 0,0475682 | 0,00361658 |
| 676 | 0,140681 | 1,15945 | 0,0186536 | 0,0455641 | 0,00617531 |
| 677 | 0,13922 | 1,13102 | 0,0359742 | 0,0441179 | 0,00785486 |
| 678 | 0,118093 | 1,08384 | 0,0726437 | 0,0439679 | 0,00749192 |
| 679 | 0,113963 | 1,0507 | 0,0777503 | 0,0433295 | 0,0076756 |
| 680 | 0,124524 | 1,02968 | 0,0555781 | 0,0422689 | 0,0083317 |
| 681 | 0,121036 | 0,999041 | 0,0344483 | 0,0411793 | 0,00890162 |
| 682 | 0,113799 | 0,965824 | 0,0143608 | 0,0400895 | 0,00944805 |
| 683 | 0,119052 | 0,941033 | 0,02674 | 0,0393597 | 0,01005 |
| 684 | 0,116364 | 0,910669 | 0,0358455 | 0,0386786 | 0,0100215 |
| 685 | 0,092509 | 0,865454 | 0,0362268 | 0,0381276 | 0,00831311 |
| 686 | 0,084993 | 0,835331 | 0,0366328 | 0,0368987 | 0,00913696 |
| 687 | 0,0920624 | 0,818681 | 0,0370609 | 0,0350647 | 0,0122213 |
| 688 | 0,0913889 | 0,794286 | 0,0540286 | 0,0353563 | 0,0114126 |
| 689 | 0,0879575 | 0,767529 | 0,0732947 | 0,0360175 | 0,00978241 |
| 690 | 0,0726076 | 0,735747 | 0,0555429 | 0,0332111 | 0,0119866 |
| 691 | 0,0654174 | 0,71133 | 0,0401845 | 0,0317476 | 0,0129781 |
| 692 | 0,0751812 | 0,702219 | 0,0297989 | 0,0330741 | 0,0114499 |
| 693 | 0,0786743 | 0,676538 | 0,0121934 | 0,0337828 | 0,011196 |

| | | | | | |
|---|---|---|---|---|---|
| 694 | 0,0779951 | 0,639831 | 0,0102158 | 0,0340804 | 0,01179 |
| 695 | 0,0671071 | 0,613639 | 0,00253648 | 0,0328576 | 0,0126253 |
| 696 | 0,0630395 | 0,591369 | 0,0161534 | 0,0319994 | 0,0127907 |
| 697 | 0,0999608 | 0,581603 | 0,0103621 | 0,0339896 | 0,00944437 |
| 698 | 0,106415 | 0,557986 | 0,0216958 | 0,0340057 | 0,00903175 |
| 699 | 0,0767181 | 0,517931 | 0,0533502 | 0,0316791 | 0,0121003 |
| 700 | 0,0703148 | 0,515768 | 0,0399009 | 0,0305392 | 0,012716 |
| 701 | 0,0722484 | 0,523714 | 0,0108522 | 0,0297844 | 0,0125804 |
| 702 | 0,0864509 | 0,490842 | 0,0323835 | 0,0289558 | 0,0133814 |
| 703 | 0,092221 | 0,465452 | 0,0505105 | 0,0283236 | 0,0134169 |
| 704 | 0,0829691 | 0,453392 | 0,0239081 | 0,0280412 | 0,0120887 |
| 705 | 0,0733356 | 0,439113 | 0,0223611 | 0,0261486 | 0,0148192 |
| 706 | 0,0635145 | 0,423742 | 0,0331353 | 0,0234642 | 0,0195456 |
| 707 | 0,0448635 | 0,38801 | 0,0301814 | 0,0241209 | 0,016491 |
| 708 | 0,032031 | 0,36479 | 0,0178392 | 0,0247937 | 0,0139962 |
| 709 | 0,0326009 | 0,370436 | 0,0152471 | 0,0253974 | 0,0130089 |
| 710 | 0,0484109 | 0,343832 | 0,0252562 | 0,0235364 | 0,0152991 |
| 711 | 0,0730509 | 0,298544 | 0,0218941 | 0,0202474 | 0,0194884 |
| 712 | 0,0634132 | 0,311893 | 0,0490264 | 0,0208898 | 0,0170493 |
| 713 | 0,0532576 | 0,31342 | 0,0605588 | 0,021251 | 0,0156937 |
| 714 | 0,0448607 | 0,282331 | 0,0331091 | 0,020617 | 0,0174354 |
| 715 | 0,0424985 | 0,253103 | 0,0235989 | 0,0196172 | 0,0180905 |
| 716 | 0,0435086 | 0,224915 | 0,0241135 | 0,0184131 | 0,0181385 |
| 717 | 0,0652168 | 0,237742 | 0,00260979 | 0,0147872 | 0,0230044 |
| 718 | 0,0742273 | 0,233963 | 0,00242534 | 0,0133886 | 0,0247997 |
| 719 | 0,0579929 | 0,196832 | 0,0251651 | 0,016389 | 0,0204948 |
| 720 | 0,0225477 | 0,159515 | 0,0173133 | 0,0182287 | 0,0191201 |
| 721 | 0,0157174 | 0,124742 | 0,0052929 | 0,0194944 | 0,0189966 |
| 722 | 0,038411 | 0,133356 | 0,0153377 | 0,0196693 | 0,0183554 |
| 723 | 0,0565938 | 0,143442 | 0,0300023 | 0,0178048 | 0,0202861 |
| 724 | 0,0258724 | 0,155529 | 0,0509521 | 0,0131656 | 0,0257159 |
| 725 | 0,00678529 | 0,131741 | 0,0812356 | 0,0130592 | 0,0250458 |
| 726 | 0,000787159 | 0,100112 | 0,0992127 | 0,0133299 | 0,0234661 |
| 727 | 0,0433495 | 0,0751799 | 0,0464366 | 0,00980402 | 0,0250624 |
| 728 | 0,0799897 | 0,0589973 | 0,00442298 | 0,0105163 | 0,023998 |
| 729 | 0,111862 | 0,0498592 | 0,0289256 | 0,0146409 | 0,0207915 |
| 730 | 0,0646497 | 0,023477 | 0,00131929 | 0,0119233 | 0,0235854 |
| 731 | 0,0363189 | 0,0199325 | 0,0243063 | 0,0107782 | 0,0249366 |
| 732 | 0,0557337 | 0,0642869 | 0,0247699 | 0,0136412 | 0,0226448 |
| 733 | 0,0442946 | 0,0441071 | 0,0298099 | 0,0124287 | 0,0253048 |
| 734 | 0,0279315 | 0,00806852 | 0,0349032 | 0,00984274 | 0,0290494 |
| 735 | 0,0539221 | 0,0242396 | 0,0292841 | 0,00812069 | 0,0279215 |
| 736 | 0,069675 | 0,00981569 | 0,0161131 | 0,00698364 | 0,03013 |
| 737 | 0,0767067 | 0,0306706 | 0,00349101 | 0,00634494 | 0,0351805 |
| 738 | 0,0424161 | 0,0415213 | 0,022353 | 0,0102875 | 0,0280115 |

| | | | | | |
|---|---|---|---|---|---|
| 739 | 0,0197773 | 0,0444898 | 0,0322521 | 0,011162 | 0,0247292 |
| 740 | 0,0204597 | 0,0365616 | 0,0111781 | 0,00642923 | 0,0290938 |
| 741 | 0,0545605 | 0,061783 | 0,0191865 | 0,00754608 | 0,0271532 |
| 742 | 0,0787423 | 0,0853639 | 0,0617704 | 0,0101292 | 0,0241748 |
| 743 | 0,0372112 | 0,0726633 | 0,199487 | 0,0124996 | 0,023339 |
| 744 | 0,0532058 | 0,102777 | 0,179677 | 0,00897573 | 0,0292764 |
| 745 | 0,093054 | 0,148505 | 0,0862467 | 0,00270963 | 0,0386471 |
| 746 | 0,0682552 | 0,125066 | 0,088186 | 0,000136095 | 0,0465496 |
| 747 | 0,0838766 | 0,0976296 | 0,0659618 | 0,00235331 | 0,0451285 |
| 748 | 0,134538 | 0,0667286 | 0,0227904 | 0,00872357 | 0,0356249 |
| 749 | 0,0235556 | 0,0279428 | 0,123724 | 0,0103132 | 0,0369584 |
| 750 | 0,060505 | 0,0373203 | 0,145307 | 0,0107538 | 0,0391939 |
| 751 | 0,106663 | 0,108698 | 0,0620343 | 0,00981794 | 0,0423636 |
| 752 | 0,043339 | 0,0686252 | 0,00041864 | 0,00827124 | 0,0453605 |
| 753 | 0,0111355 | 0,0674569 | 0,0075459 | 0,00714666 | 0,0460654 |
| 754 | 0,0275628 | 0,158013 | 0,0751667 | 0,00689508 | 0,0427425 |
| 755 | 0,0309862 | 0,19917 | 0,163345 | 0,00388474 | 0,0478053 |
| 756 | 0,0182781 | 0,216233 | 0,21352 | 0,00210889 | 0,0517013 |
| 757 | 0,0252317 | 0,201353 | 0,174196 | 0,00445095 | 0,0489948 |
| 758 | 0,00827941 | 0,191166 | 0,132089 | 0,00525887 | 0,0508289 |
| 759 | 0,0565786 | 0,192297 | 0,118286 | 0,00504109 | 0,0542565 |
| 760 | 0,0872427 | 0,22075 | 0,183724 | 0,00318165 | 0,0586112 |
| 761 | 0,0658363 | 0,300066 | 0,104753 | 0,0100471 | 0,0519982 |
| 762 | 0,0241394 | 0,376482 | 0,0184952 | 0,0171744 | 0,0438133 |
| 763 | 0,0284383 | 0,396572 | 0,0464148 | 0,0168777 | 0,0412992 |
| 764 | 0,000124866 | 0,303627 | 0,130614 | 0,00616139 | 0,0545737 |
| 765 | 0,00461788 | 0,213718 | 0,196409 | 0,000926733 | 0,0639809 |
| 766 | 0,13554 | 0,238123 | 0,156985 | 0,0120965 | 0,0475753 |
| 767 | 0,247238 | 0,293711 | 0,138956 | 0,0105002 | 0,056061 |
| 768 | 0,290377 | 0,348456 | 0,094015 | 0,00847673 | 0,0644308 |
| 769 | 0,183399 | 0,377787 | 0,0341969 | 0,0163129 | 0,0539056 |
| 770 | 0,101286 | 0,280839 | 0,119374 | 0,0117435 | 0,0567976 |
| 771 | 0,0601417 | 0,244812 | 0,26295 | 0,00796885 | 0,0618305 |
| 772 | 0,0890743 | 0,391197 | 0,251532 | 0,0117279 | 0,0647202 |
| 773 | 0,00777727 | 0,37023 | 0,207107 | 0,0105413 | 0,0625246 |
| 774 | 0,0757178 | 0,321828 | 0,148411 | 0,0155419 | 0,055105 |
| 775 | 0,136806 | 0,271536 | 0,0761395 | 0,0309141 | 0,0410861 |
| 776 | 0,0701587 | 0,228801 | 0,0737386 | 0,0191902 | 0,0556996 |
| 777 | 0,11258 | 0,222741 | 0,126822 | 0,0175435 | 0,0572481 |
| 778 | 0,265515 | 0,252602 | 0,221183 | 0,0276195 | 0,0443252 |
| 779 | 0,188333 | 0,246527 | 0,0535802 | 0,0288538 | 0,050003 |
| 780 | 0,157566 | 0,244106 | 0,0132321 | 0,0293459 | 0,0522663 |

**Table S2.** Spectral power distribution raw data. Irradiance (mW/m$^2$) in each light condition at 1 nanometer (nm) intervals from 380 to 780 nm. BWL = Bright White Light. DL = Dim Light. 'DL (washout') refers to the 1-hour exposure of warm-white DL preceding light adaptation. LWL = Long-Wavelength Light. SWL = Short-Wavelength Light.

### Supplementary figures, tables, and results from exploratory statistical analyses

#### RTs trial-by-trial on the PVT in each light condition, stratified by measurement block 1-3

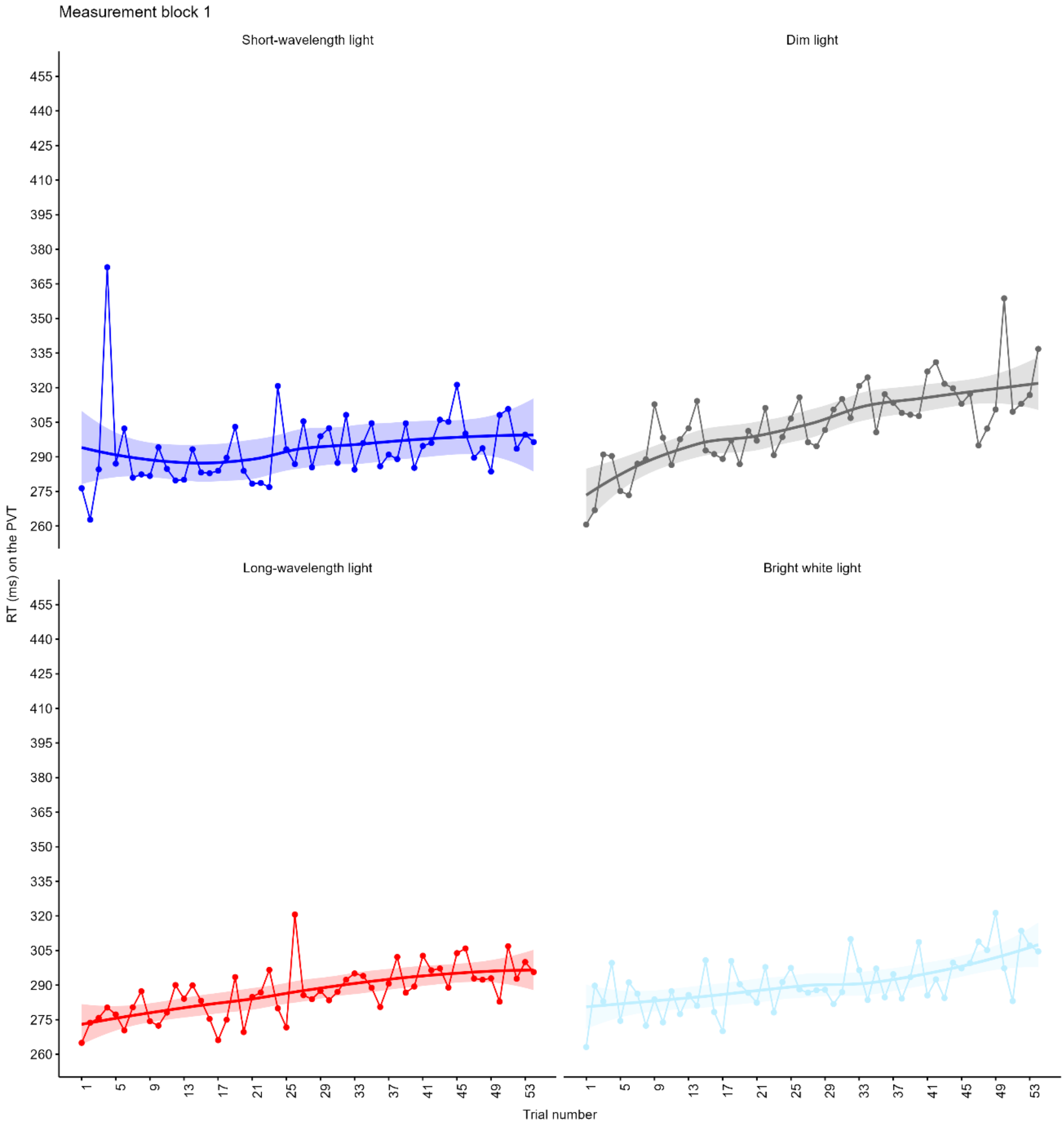


**Figure S3.** Line plot of RTs (averaged across the study sample of 39 subjects) from trial 1 to 54 on the PVT in each light condition during the 1st measurement block. The smoothing line represents 'loess' smoothing with 95% confidence interval as error band.

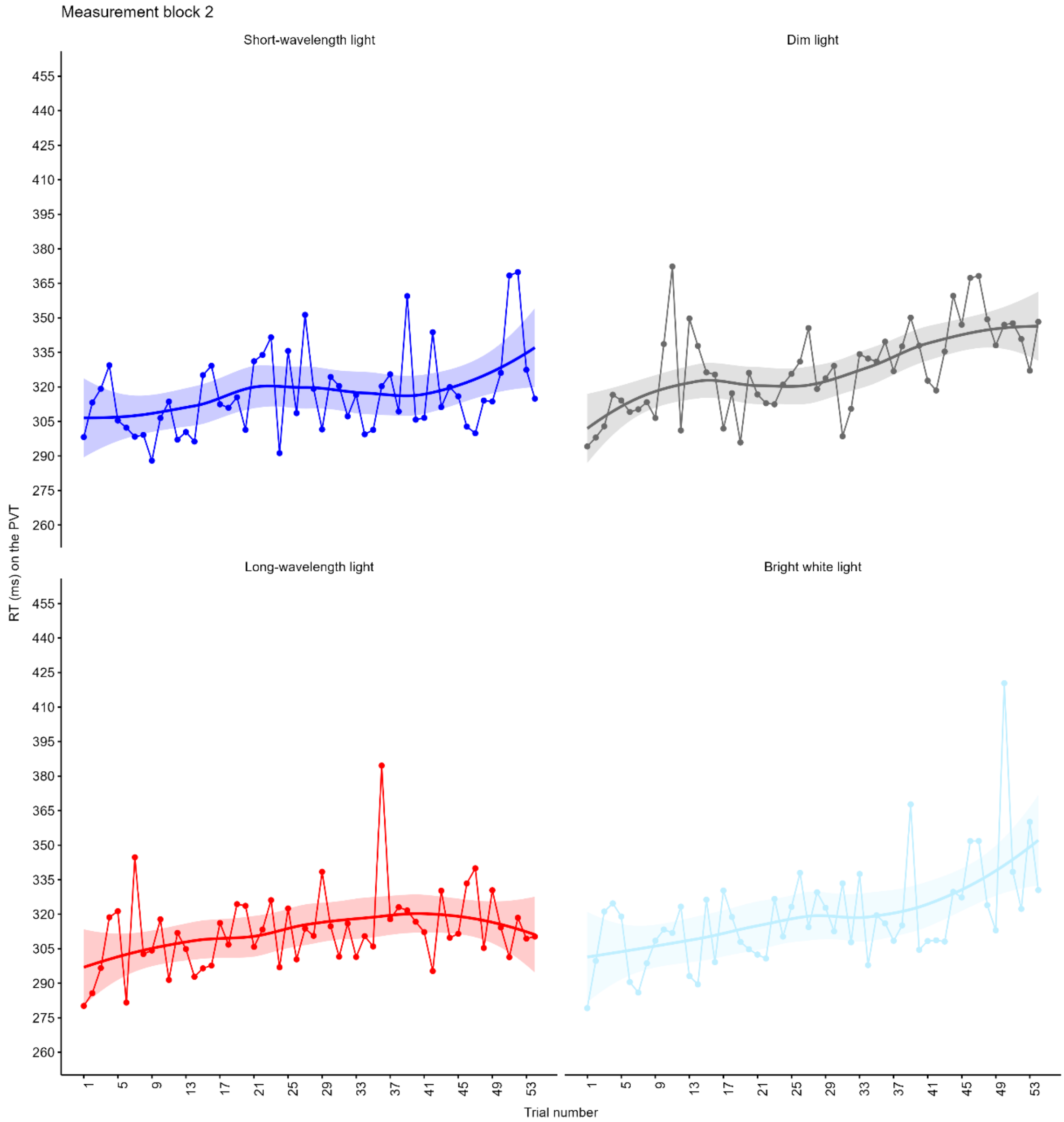


**Figure S4.** Line plot of RTs (averaged across the study sample of 39 subjects) from trial 1 to 54 on the PVT in each light condition during the 2$^{nd}$ measurement block. The smoothing line represents 'loess' smoothing with 95% confidence interval as error band.

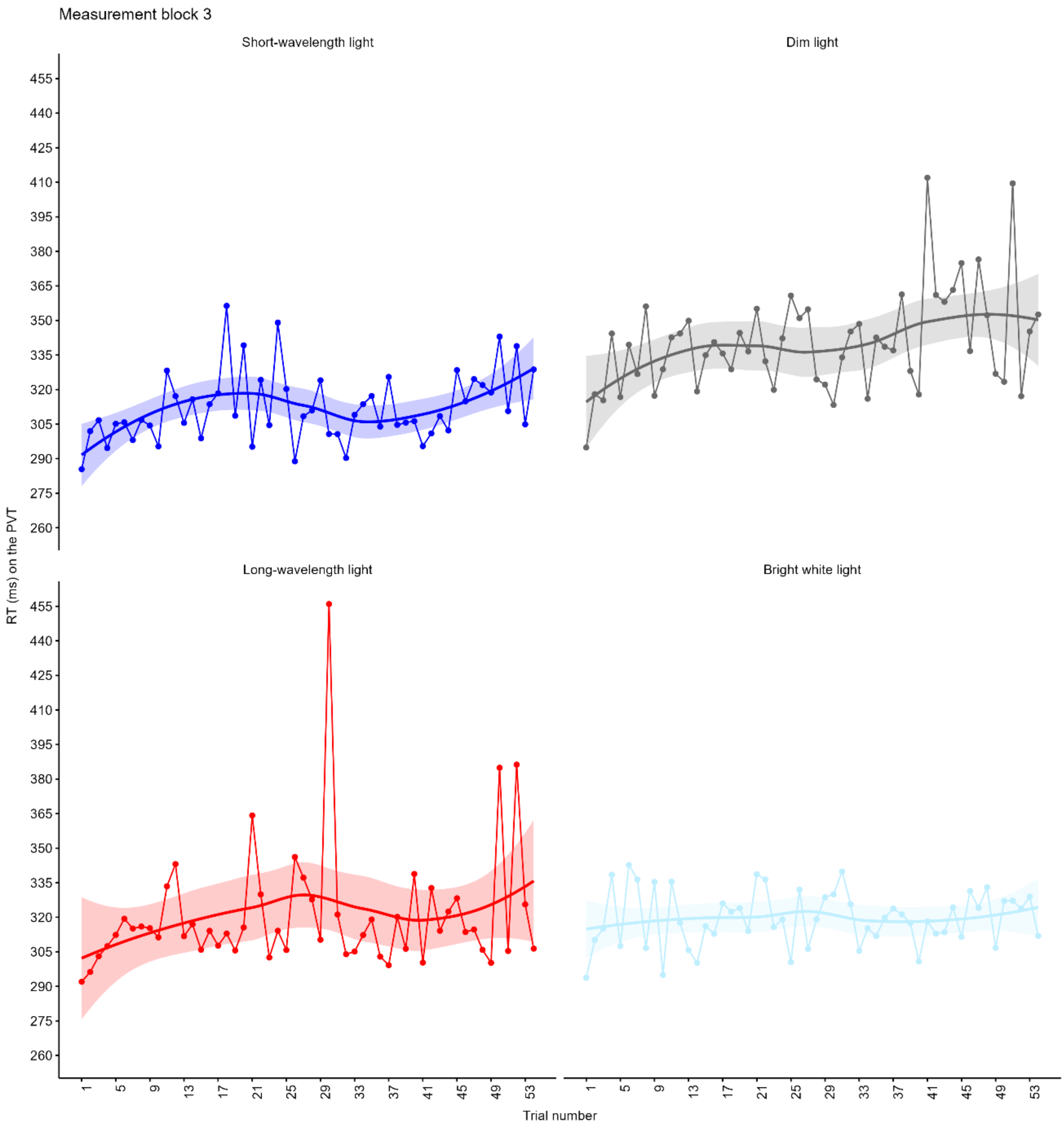


**Figure S5.** Line plot of RTs (averaged across the study sample of 39 subjects) from trial 1 to 54 on the PVT in each light condition during the 3rd measurement block. The smoothing line represents 'loess' smoothing with 95% confidence interval as error band.

### Lapses

#### *Main Effects Model*

| | Lapses | | | | |
|---|---|---|---|---|---|
| *Parameters* | *IRR (95% CI)* | *SE* | *z* | *p* | $r_p$ |
| Intercept | 1.04 (0.19 – 5.68) | 0.90 | 0.50 | 0.562 | |
| **Light** | | | | | |
| SWL | 0.60 (0.42 – 0.85) | 0.11 | -2.87 | **.004** | -0.42 |
| LWL | 0.57 (0.40 – 0.82) | 0.10 | -3.10 | **.002** | -0.44 |
| BWL | 0.70 (0.49 – 0.99) | 0.12 | -2.01 | **.044** | -0.31 |
| **Block** | | | | | |
| Block 2 | 2.41 (1.82 – 3.19) | 0.34 | 6.18 | **< .001** | 0.70 |
| Block 3 | 2.48 (1.89 – 3.26) | 0.35 | 6.50 | **< .001** | 0.72 |
| **Chronotype** | | | | | |
| Eveningness | 1.09 (0.68 – 1.75) | 0.26 | 0.37 | .714 | 0.06 |
| Morningness | 1.07 (0.60 – 1.91) | 0.32 | 0.22 | .830 | 0.04 |
| **Season** | | | | | |
| Spring | 1.92 (1.08 – 3.44) | 0.57 | 2.21 | **.027** | 0.33 |
| Summer | 1.38 (0.68 – 2.81) | 0.50 | 0.89 | .373 | 0.14 |
| Winter | 0.97 (0.97 – 1.01) | 0.35 | -0.08 | .934 | -0.01 |
| Sleep efficiency | 0.99 (0.97 - 1.01) | 0.01 | -0.64 | .519 | -0.10 |
| **Random effects** | *Coefficient* | | | | |
| SD (Residual) | 1.29 | | | | |
| SD (Intercept: Subject) | 0.56 | | | | |
| SD (Intercept: Subject:Light) | 0.47 | | | | |
| SD (Intercept: Subject:Block) | 0.22 | | | | |
| Adjusted / Unadjusted ICC | 0.396 / 0.330 | | | | |
| Conditional / Marginal $R^2$ | 0.497 / 0.168 | | | | |

**Table S3.** Generalized Linear Mixed-Effects Model analyses of the number of lapses on the Psychomotor Vigilance Task. Incidence Rate Ratios (IRR) represent the exponentiated model coefficients. Significant *p*-values are highlighted in bold. BWL = Bright White Light. CI = Confidence Interval. ICC = Intra-Class Correlation Coefficient. LWL = Long-Wavelength Light. SD = Standard Deviation. SE = Standard Error. SWL = Short-Wavelength Light.

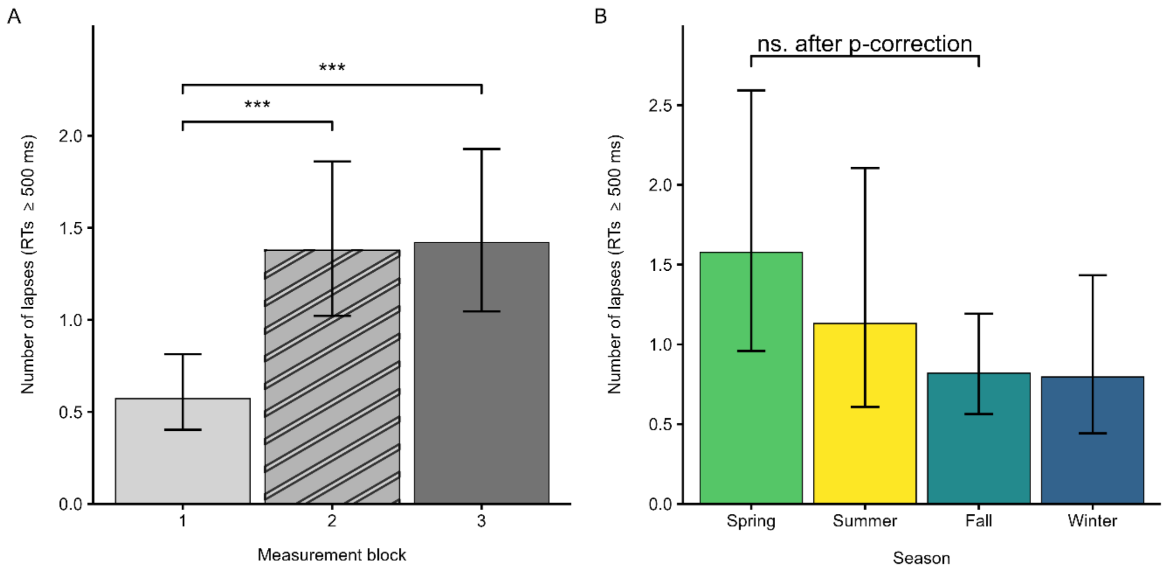


**Figure S6.** Estimated Marginal Means (EMMs) of the number of lapses across measurement blocks (A) and seasons (B). Error bars = 95% Confidence Intervals (CIs). EMMs and CIs were back-transformed from log to original scale for visualization purposes. ms = milliseconds. RTs = Reaction Times. ****p* (corrected) < .001.

### Mean Reaction Time (RT)

#### *Main Effects Model*

| | Mean RT | | | | |
|---|---|---|---|---|---|
| *Parameters* | *Coefficient (95% CI)* | *SE* | *t*(422) | *p* | $f^2$ |
| Intercept | 291.97 (250.83 – 339.85) | 22.56 | 73.47 | < .001 | |
| **Light** | | | | | |
| SWL | 0.95 (0.93 - 0.98) | 0.01 | -3.27 | **.001** | 0.11 |
| LWL | 0.96 (0.93 - 0.99) | 0.01 | -2.99 | **.003** | 0.09 |
| BWL | 0.97 (0.94 – 1.00) | 0.01 | -2.10 | **.036** | 0.04 |
| **Block** | | | | | |
| Block 2 | 1.08 (1.06 - 1.10) | 9.20e-03 | 9.10 | **< .001** | 1.10 |
| Block 3 | 1.10 (1.08 - 1.12) | 9.34e-03 | 10.95 | **< .001** | 1.60 |
| **Chronotype** | | | | | |
| Eveningness | 1.02 (0.97 - 1.06) | 0.02 | 0.75 | .452 | e.78e-03 |
| Morningness | 1.01 (0.96 - 1.06) | 0.03 | 0.30 | .764 | 5.44e-04 |
| **Season** | | | | | |
| Spring | 1.04 (0.97 - 1.12) | 0.04 | 1.24 | .285 | 0.04 |
| Summer | 1.09 (1.00 - 1.19) | 0.05 | 1.92 | .064 | 0.11 |
| Winter | 0.99 (0.91 - 1.07) | 0.04 | -0.30 | .732 | 2.73e-03 |
| Sleep Efficiency | 1.00 (1.00 - 1.00) | 0.00 | -0.09 | .363 | 4.24e-05 |
| **Random Effects** | | *Coefficient* | | | |
| SD (Residual) | | 0.07 | | | |
| SD (Intercept: Subject) | | 0.08 | | | |
| SD (Intercept: Subject:Light) | | 0.05 | | | |
| SD (Intercept: Subject:Block) | | 8.70e-03 | | | |
| Adjusted / Unadjusted ICC | | 0.633 / 0.516 | | | |
| Conditional / Marginal $R^2$ | | 0.701 / 0.185 | | | |

**Table S4**. Linear Mixed-Effects Model analyses of mean reaction time on the Psychomotor Vigilance Task. Significant *p*-values are highlighted in bold. Model coefficients are exponentiated, as the models provide coefficients on the log-scale. BWL = Bright White Light. CI = Confidence Interval. ICC = Intra-Class Correlation Coefficient. LWL = Long-Wavelength Light. RT = Reaction Time. SD = Standard Deviation. SE = Standard Error. SWL = Short-Wavelength Light.

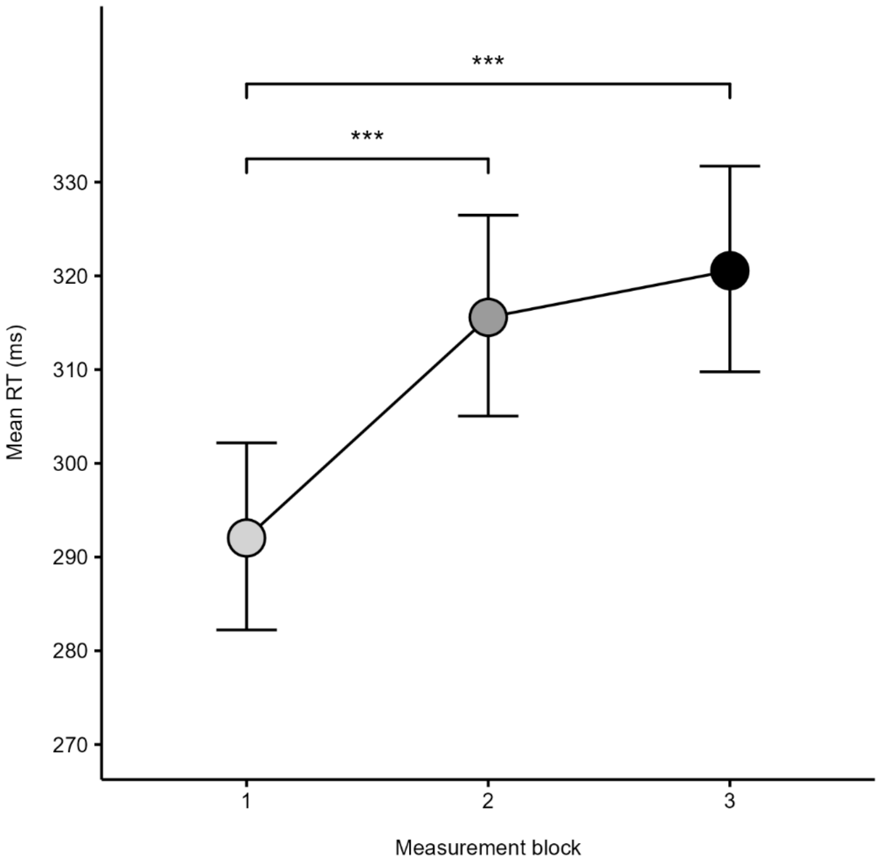


**Figure S7.** Estimated Marginal Means (EMMs) of mean Reaction Time (RT) across measurement blocks. The y-axis limit has been adjusted so that group differences appear more clearly. Error bars = 95% confidence intervals (CIs). EMMs and CIs were back-transformed from log to original scale for visualization purposes. ms = milliseconds. ****p* (corrected) < .001.

### Reaction Time Variability (RTV)

#### *Main Effects Model*

| | RTV | | | | |
|---|---|---|---|---|---|
| *Parameters* | *Coefficient (95% CI)* | *SE* | *z* | *p* | $r_p$ |
| Intercept | 2.04e-8 (0.00 - 0.00) | 2.40E-08 | -17.67 | < .001 | |
| Mean RT | 46.99 (33.79 - 65.35) | 8.24 | 22.76 | **< .001** | 0.96 |
| **Light** | | | | | |
| SWL | 1.07 (0.99 - 1.16) | 0.04 | 1.75 | .081 | 0.27 |
| LWL | 1.07 (0.99 - 1.15) | 0.04 | 1.63 | .104 | 0.25 |
| BWL | 1.03 (0.95 - 1.11) | 0.04 | 0.75 | .452 | 0.12 |
| **Block** | | | | | |
| Block 2 | 0.87 (0.81 - 0.93) | 0.03 | -3.75 | **< .001** | -0.51 |
| Block 3 | 0.83 (0.78 - 0.89) | 0.03 | -4.79 | **< .001** | -0.61 |
| **Chronotype** | | | | | |
| Eveningness | 0.94 (0.83 - 1.06) | 0.06 | -1.19 | .235 | -0.19 |
| Morningness | 0.92 (0.79 - 1.06) | 0.07 | -1.19 | .234 | -0.19 |
| **Season** | | | | | |
| Spring | 1.08 (0.92 - 1.26) | 0.09 | 0.87 | .384 | 0.14 |
| Summer | 0.84 (0.69 - 1.03) | 0.09 | -1.71 | .087 | -0.26 |
| Winter | 1.09 (0.91 - 1.31) | 0.10 | 0.96 | .338 | 0.15 |
| Sleep efficiency | 1.00 (0.99 - 1.00) | 2.42e-03 | -1.14 | .253 | -0.18 |
| **Random effects** | *Coefficient* | | | | |
| SD (Residual) | 0.26 | | | | |
| SD (Intercept: Subject) | 0.17 | | | | |
| SD (Intercept: Subject:Light) | 0.08 | | | | |
| SD (Intercept: Subject:Block) | 0.08 | | | | |
| Adjusted / Unadjusted ICC | 0.381 / 0.130 | | | | |
| Conditional / Marginal $R^2$ | 0.789 / 0.660 | | | | |

**Table S5**. Generalized Linear Mixed-Effects Model analyses of intra-individual reaction time variability on the Psychomotor Vigilance Task. Point estimates are the exponentiated model coefficients. Significant p-values are highlighted in bold. BWL = Bright White Light. CI = Confidence Interval. ICC = Intra-Class Correlation Coefficient. LWL = Long-Wavelength Light. RT = Reaction Time. RTV = Reaction Time Variability. SD = Standard Deviation. SE = Standard Error. SWL = Short-Wavelength Light.

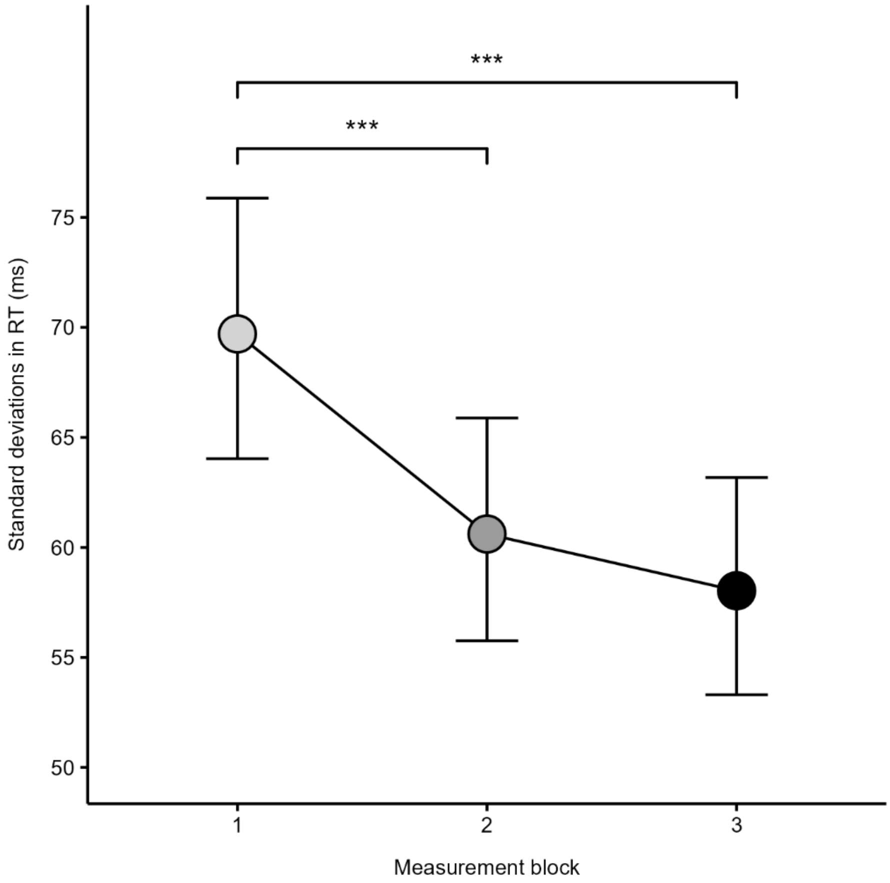


**Figure S8.** Estimated Marginal Means (EMMs) of Reaction Time (RT) Variability (standard deviations in RT) across measurement blocks. The y-axis limit has been adjusted so that group differences appear more clearly. Error bars = 95% Confidence Intervals (CIs). EMMs and CIs were back-transformed from log to original scale for visualization purposes. ms = milliseconds. ****p* (corrected) < .001.

### Mean 10% fastest RTs

#### *Main Effects Model*

| | Mean 10% Fastest RTs | | | | |
|---|---|---|---|---|---|
| *Parameters* | *Coefficient (95% CI)* | *SE* | *t*(422) | *p* | $f^2$ |
| Intercept | 241.40 (220.52 - 264.27) | 11.11 | 119.17 | < .001 | |
| **Light** | | | | | |
| SWL | 0.98 (0.97 - 1.00) | 7.76e-03 | -2.39 | **.017** | 0.06 |
| LWL | 0.98 (0.96 - 0.99) | 7.75e-03 | -2.81 | **.005** | 0.08 |
| BWL | 0.99 (0.97 - 1.00) | 7.98e-03 | -1.71 | .088 | 0.03 |
| **Block** | | | | | |
| Block 2 | 1.04 (1.03 - 1.05) | 6.68e-03 | 6.16 | **< .001** | 0.51 |
| Block 3 | 1.05 (1.03 - 1.06) | 6.71e-03 | 7.12 | **< .001** | 0.68 |
| **Chronotype** | | | | | |
| Eveningness | 1.04 (1.01 - 1.07) | 0.02 | 2.35 | **.019** | 0.03 |
| Morningness | 1.04 (1.01 - 1.08) | 0.02 | 2.29 | **.023** | 0.02 |
| **Season** | | | | | |
| Spring | 1.01 (0.96 - 1.06) | 0.03 | 0.42 | 0.676 | 4.82e-03 |
| Summer | 1.08 (1.01 - 1.16) | 0.04 | 2.37 | **.018** | 0.16 |
| Winter | 0.97 (0.92 - 1.03) | 0.03 | -0.93 | .355 | 0.03 |
| Sleep Efficiency | 1.00 (1.00 - 1.00) | 4.99e-04 | -0.96 | .337 | 5.76e-03 |
| **Random Effects** | | *Coefficient* | | | |
| SD (Residual) | | 0.05 | | | |
| SD (Intercept: Subject) | | 0.06 | | | |
| SD (Intercept: Subject:Light) | | 0.02 | | | |
| SD (Intercept: Subject:Block) | | 0.02 | | | |
| Adjusted / Unadjusted ICC | | 0.685 / 0.534 | | | |
| Conditional / Marginal $R^2$ | | 0.754 / 0.220 | | | |

**Table S6.** Linear Mixed-Effects Model analyses of the mean 10% fastest reaction times on the Psychomotor Vigilance Task. Significant *p*-values are highlighted in bold. Model coefficients are exponentiated, as the models provide coefficients on the log-scale. BWL = Bright White Light. CI = Confidence Interval. ICC = Intra-Class Correlation Coefficient. LWL = Long-Wavelength Light. RT = Reaction Time. SD = Standard Deviation. SE = Standard Error. SWL = Short-Wavelength Light.

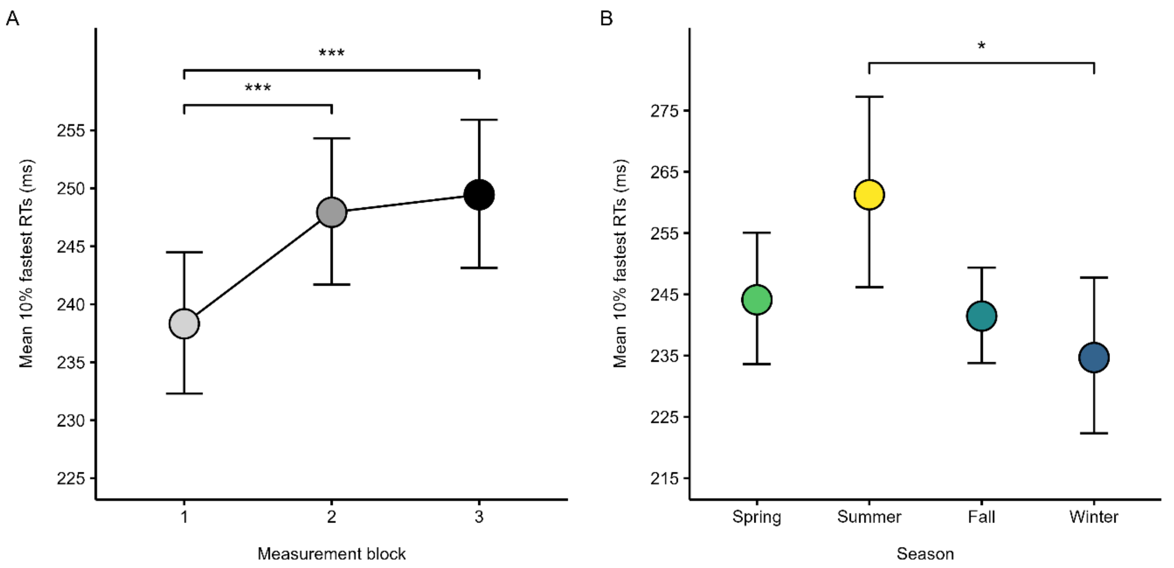


**Figure S9.** Estimated Marginal Means (EMMs) of the mean 10% fastest Reaction Times (RTs) across measurement blocks (A) and seasons (B). Y-axis limits have been adjusted so that group differences appear more clearly. Error bars = 95% Confidence Intervals (CIs). EMMs and CIs were back-transformed from log to original scale for visualization purposes. ms = milliseconds. ***$p$ (corrected) < .001.

### Results from complete-case analyses

The same set of statistical analyses reported in the paper was performed on a dataset with complete-cases (listwise deletion, resulting in N = 33) to assess the sensitivity of the imputed values. Overall, the pattern of results was similar to the results from analyses on imputed data, except for a significant interaction between light and chronotype on lapses (DL eveningness*SWL intermediate type, $p_{corrected}$ = .048, Fig. S11) and a significant main effect of light on RTV (Fig. S13a). However, these findings should be interpreted cautiously, as estimates are often biased (smaller standard errors and p-values, narrower confidence intervals) in a complete-case analysis when missing data are not missing completely at random (MCAR)[3,4]. Across all PVT outcome measures, there were no significant interactions between light and block, no significant main effect of sleep efficiency, and no sex differences in performance.

#### Lapses

The initial model with the interaction term light*block revealed significant simple effects of light and block but no significant interactions, hence, light and block were modelled as main effects in a new model.

##### *Main Effects Model*

Analysis of the number of lapses showed significant main effects of light (short-wavelength light [SWL]: $p$ < .001; long-wavelength light [LWL]: $p$ = .007; bright white light [BWL]: $p$ = .037) and block (block 2 and 3: $p$'s < .001). Post-hoc analyses on Estimated Marginal Means (EMMs) with Tukey's Honestly Significant Difference (HSD) test correcting for multiple comparisons showed that subjects had significantly fewer lapses in SWL ($p$ = .002 *EMM* = 0.88[0.59-1.31], *SE* = 1.23) and LWL ($p$ = .037, *EMM* = 1.04[0.69-1.55], *SE* = 1.23) compared to dim light (DL, *EMM* = 1.70[1.15-2.51], *SE* = 1.22), see Fig. 12a. Post-hoc analyses on the main effect of block showed that subjects displayed a significantly higher number of lapses in block 2 ($p < .001$, *EMM* = 1.55[1.10-2.20], *SE* = 1.19) and 3 ($p < .001$, *EMM* = 1.57[1.11-2.23], *SE* = 1.20), relative to block 1 (*EMM* = 0.64[0.43-0.95], *SE* = 1.23) (Fig. S12b). There was also a significant main effect of season (spring: $p$ = .038), but the effect was no longer significant after correcting for multiple comparisons.

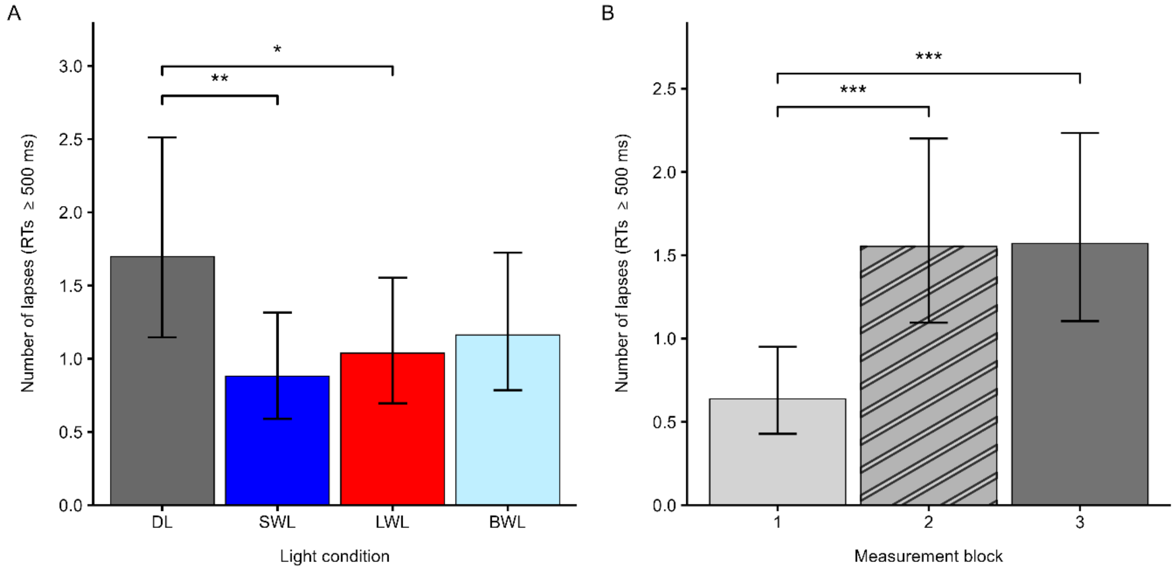


**Figure S10.** Estimated Marginal Means (EMMs) of the number of lapses in each light condition (A) and across measurement blocks (B). Error bars = 95% Confidence Intervals (CIs). EMMs and CIs were back-transformed from log to original scale for visualization purposes. BWL = Bright White Light. DL = Dim Light. LWL = Long-Wavelength Light. ms = milliseconds. RT = Reaction Time. SWL = Short-Wavelength Light. ****p* (corrected) < .001. ***p* (corrected) < .01. **p* (corrected) < .05.

#### ***Models with Interaction Terms (Light x Season, Light x Chronotype)***:

There was no significant interaction effect between light and season, however, light interacted significantly with chronotype (BWL*morningness: $p$ = .021). Post-hoc analyses did not reveal any significant differences within the same light conditions nor chronotypes, but 'evening' types tested in DL had a higher number of lapses compared to 'intermediate' types tested in SWL ($p$ = .048, Fig. S13).

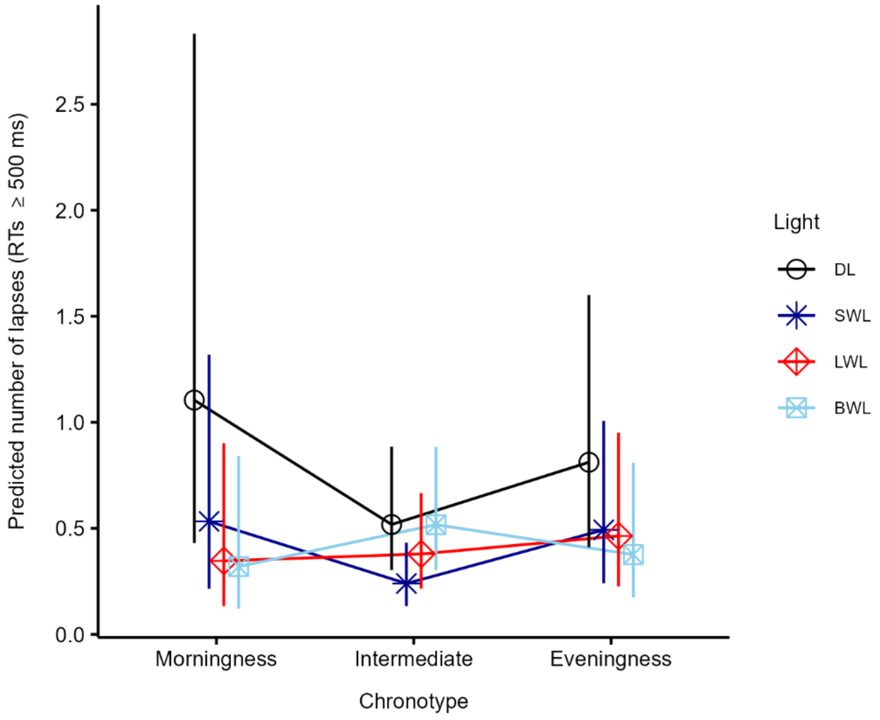


**Figure S11.** Predicted number of lapses as a function of the interaction between light and chronotype. Error bars represent 95% Confidence Intervals. BWL = Bright White Light. DL = Dim Light. LWL = Long-Wavelength Light. ms = milliseconds. RT = Reaction Time. SWL = Short-Wavelength Light. Significant difference: DL – eveningness vs. SWL – intermediate type (*p* (corrected) = .048).

### Mean RT

#### *Main Effects Model*

There were significant main effects of light conditions (SWL: $p$ < .001; LWL: $p$ = .003; BWL: $p$ = .042) and block (block 2 and 3, $p$'s < .001). Post-hoc analyses of EMMs showed that participants responded significantly faster in SWL ($p$ = .003, *EMM* = 307.85[294.87-321.39], *SE* = 1.02) and LWL ($p$ = .019, *EMM* = 310.42[297.19-324.25], *SE* = 1.02) compared to DL (*EMM* = 325.91[311.70-340.76], *SE* = 1.02), see Fig. S14a. However, they responded slower over time in testing; mean RT was slower in block 2 ($p$ < .001, *EMM* = 320.36[307.43-333.84], *SE* = 1.02) and 3 ($p$ < .001, *EMM* = 326.69[313.52-340.42], *SE* = 1.02), relative to block 1 (*EMM* = 298.06[286.04-310.59], *SE* = 1.02) (Fig. S14b).

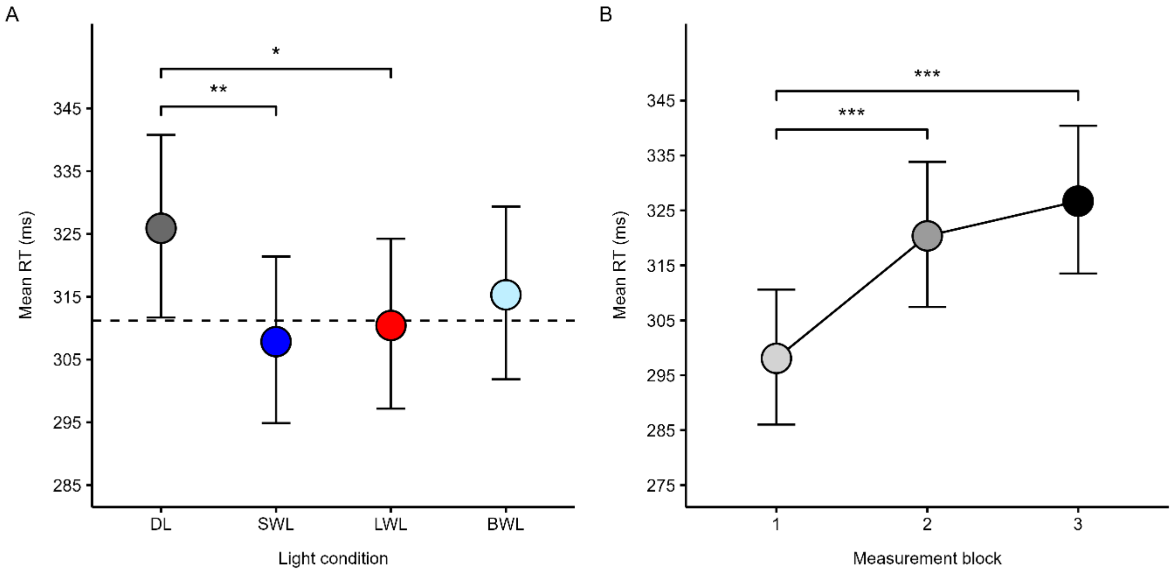


**Figure S12.** Estimated Marginal Means (EMMs) of mean Reaction Time (RT) in each light condition (A) and across measurement blocks (B). The y axis limit has been adjusted so that group differences appear more clearly. Error bars = 95% Confidence Intervals (CIs). Dashed line in panel A represents mean estimated RT across SWL, LWL and BWL. EMMs and CIs were back-transformed from log to original scale for visualization purposes. BWL = Bright White Light. DL = Dim Light. LWL = Long-Wavelength Light. ms = milliseconds. SWL = Short-Wavelength Light. ****p* (corrected) < .001. ***p* (corrected) < .01. **p* (corrected) < .05.

##### ***Models with Interaction Terms (Light x Season, Light x Chronotype)***:

There was no significant interaction between light and season. However, light interacted significantly with chronotype (LWL*morningness: $p$ = .039; BWL*morningness: $p$ = .032), but the effects were no longer significant after correcting for multiple comparisons.

#### RTV

The initial model with the interaction term light*block revealed significant simple effects of block but no significant interactions between light and block, hence, block and light were modelled as main effects in a new model, with an additional fixed effect: mean RT, as the mean and SD in RT data are often correlated to some extent[5].

##### ***Main Effects Model***

There was a significant main effect of mean RT ($p$ < .001), in that variability in RT increased with longer RTs (Pearsons' partial $r$ =.76 [95% CI: .71-.80], $p$ < .001). There was also a significant main effect of light (LWL: $p$ = .006) and block (block 2 and 3: $p$'s < .001). Post-hoc analyses on EMMs showed that subjects expressed larger variability in LWL ($p$ = .031, *EMM* = 67.45[60.87-74.74], *SE* = 1.05) compared to DL (*EMM* = 60.09[54.00-66.86], *SE* = 1.06), see Fig. S15a. However, they expressed lower variability in block 2 ($p$ < .001, *EMM* = 61.34[55.36-67.96], *SE* = 1.05) and 3 ($p$ < .001, *EMM* = 58.44[52.71-64.79], *SE* = 1.05), relative to block 1 (*EMM* = 71.85[64.82-79.65], *SE* = 1.05) (Fig. S15b). There were no significant main effects of season nor chronotype.

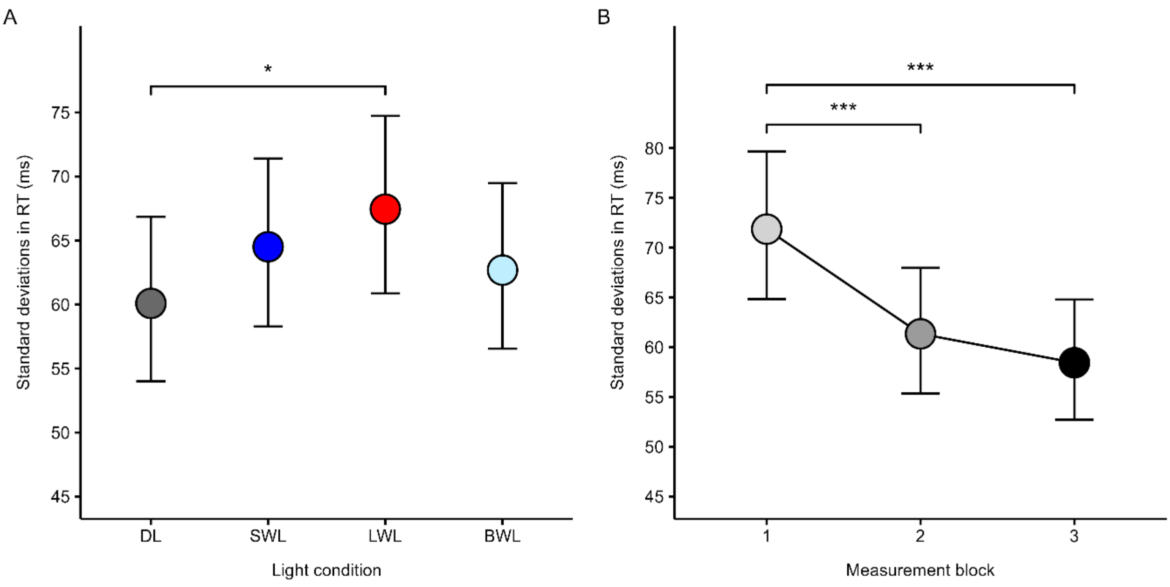


**Figure S13.** Estimated Marginal Means (EMMs) of Reaction Time (RT) Variability (standard deviations in RT) in each light condition (A) and across measurement blocks (B). Error bars = 95% Confidence Intervals (CIs). The y axis limit has been adjusted so that group differences appear more clearly. EMMs and CIs were back transformed from log to original scale for visualization purposes. BWL = Bright White Light. DL = Dim Light. LWL = Long-Wavelength Light. ms = milliseconds. SWL = Short-Wavelength Light. ****p* (corrected) < .001. **p* (corrected) < .05.

##### *Models with Interaction Terms (Light x Season, Light x Chronotype):*

There were no significant interactions between light and season nor between light and chronotype.

#### Mean of the 10% Fastest RTs

Both light and block had significant simple effects on the mean 10% fastest RTs, hence, light and block were modelled as main effects in a new model.

##### *Main Effects Model*

There were significant main effects of light conditions (SWL: $p$ = .001; LWL: $p$ = .003), block (block 2 and 3, $p$'s < .001), chronotype (eveningness: $p$ = .040), and season (summer: $p$ = .025). Post-hoc analyses of EMMs showed that subjects' mean 10% fastest RTs were lower (that is, participants responded faster) in SWL ($p$ = .007, *EMM* = 245.15[237.53-253.02], *SE* = 0.02) and LWL ($p$ = .018, *EMM* = 245.70[238.02-253.63], *SE* = 0.02) compared to DL (*EMM* = 251.87[243.91-260.09], *SE* = 0.02), see Fig. S16a. However, they responded slower over time; 10% fastest RTs were slower in block 2 ($p$ < .001, *EMM* = 250.03[242.32-257.98], *SE* = 0.02) and 3 ($p$ < .001, *EMM* = 252.66[244.88-260.69], *SE* = 0.02), relative to block 1 (*EMM* = 240.77[233.35-248.42], *SE* = 0.02) (Fig. S16b). The main effect of season and chronotype did not remain significant after correcting for multiple comparisons.

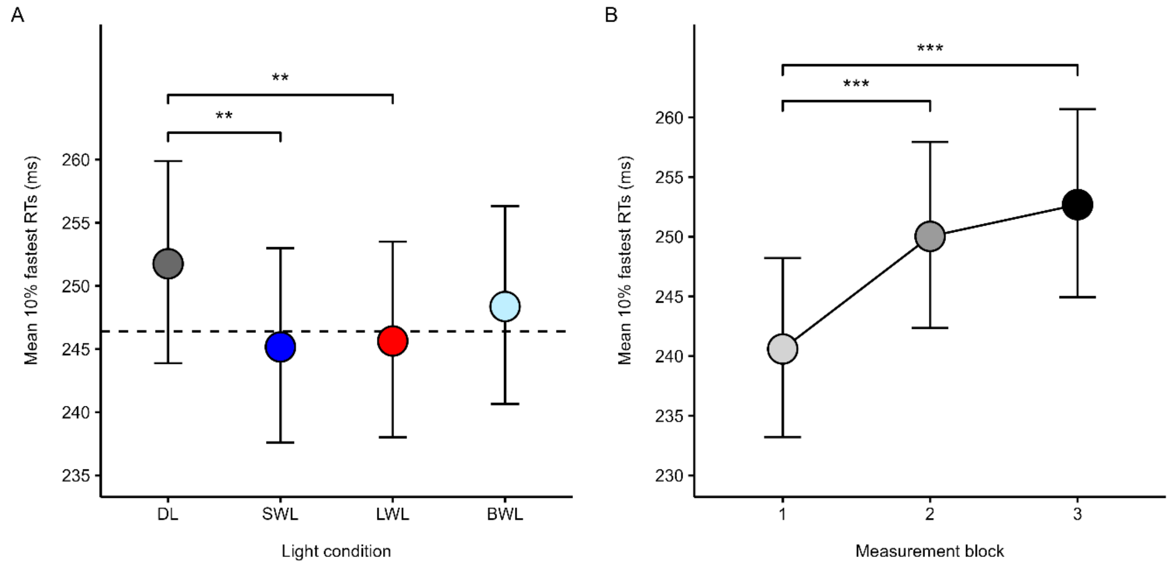


**Figure S14.** Estimated Marginal Means (EMMs) of the mean of the 10% fastest Reaction Times (RTs) in each light condition (A) and across measurement blocks (B). The y axis limit has been adjusted so that group differences appear more clearly. Error bars = 95% Confidence Intervals (CIs). Dashed line in panel A represents the mean 10% fastest RT across SWL, LWL, and BWL. EMMs and CIs were back transformed from log to original scale for visualization purposes. BWL = Bright White Light. DL = Dim Light. LWL = Long-Wavelength Light. ms = milliseconds. RT = Reaction Time. SWL = Short-Wavelength Light. ***$p$ (corrected) < .001. **$p$ (corrected) < .01.

#### ***Models with Interaction Terms (Light x Season, Light x Chronotype)***:

There was no significant interactions effect between light and season nor between light and chronotype.

### "Crude" model estimates

Below follow model estimates from mixed model analyses of PVT outcomes, unadjusted for chronotype, season, and sleep efficiency. Light and block were modeled as fixed effects, subject was modeled with random intercept, and a complex random intercept[6] was included for light and block.

| | Lapses | | | | | |
|---|---|---|---|---|---|---|
| *Parameters* | *IRR (95% CI)* | *EMM (95% CI)* | *SE* | *z* | $p_{corr}$ | $r_p$ |
| Intercept | 0.76 (0.51 – 1.13) | | 0.15 | -1.37 | | |
| **Light** | | | | | | |
| SWL | 0.62 (0.44 – 0.88) | 0.85 (0.61 – 1.20) | 0.11 | -2.70 | **.035** | -0.40 |
| LWL | 0.59 (0.42 – 0.84) | 0.82 (0.58 – 1.16) | 0.11 | -2.93 | **.019** | -0.42 |
| BWL | 0.69 (0.48 – 0.98) | 0.94 (0.67 – 1.35) | 0.12 | -2.10 | .153 | -0.32 |
| **Block** | | | | | | |
| Block 2 | 2.41 (1.83 – 3.18) | 1.30 (0.98 – 1.72) | 0.34 | 6.23 | **< .001** | 0.71 |
| Block 3 | 2.48 (1.89 – 3.25) | 1.34 (1.00 – 1.78) | 0.34 | 6.54 | **< .001** | 0.72 |
| **Random effects** | *Coefficient* | | | | | |
| SD (Residual) | 1.31 | | | | | |
| SD (Intercept: Subject) | 0.63 | | | | | |
| SD (Intercept: Subject:Light) | 0.47 | | | | | |
| SD (Intercept: Subject:Block) | 0.20 | | | | | |
| Adjusted / Unadjusted ICC | | 0.428 / 0.376 | | | | |
| Conditional / Marginal $R^2$ | | 0.499 / 0.123 | | | | |

**Table S7.** Generalized Linear Mixed-Effects Model analysis of the number of lapses on the Psychomotor Vigilance Task. Incidence Rate Ratio (IRR) values represent the exponentiated coefficients. Siginificant *p*-values corrected for multiple comparisons using Tukey's HSD are highlighted in bold. Pearson's partial *r* ($r_p$) was approximated as effect size[7]. BWL = Bright White Light. CI = Confidence Interval. EMM = Estimated Marginal Mean. ICC = Intra-Class Correlation Coefficient. LWL = Long-Wavelength Light. SD = Standard Deviation. SE = Standard Error. SWL = Short-Wavelength Light.

| | Mean RT | | | | | |
|---|---|---|---|---|---|---|
| *Parameters* | *Coefficient (95% CI)* | *EMM (95% CI)* | *SE* | *t*(428) | $p_{corr}$ | $f^2$ |
| Intercept | 297.63 (287.42 – 308.21) | | 5.29 | 320.47 | | |
| **Light** | | | | | | |
| SWL | 0.95 (0.93 – 0.98) | 300.53 (290.79 – 310.59) | 0.01 | -3.25 | **.008** | 0.10 |
| LWL | 0.96 (0.93 – 0.99) | 301.64 (291.86 – 311.74) | 0.01 | -3.00 | **.018** | 0.09 |
| BWL | 0.97 (0.94 – 1.00) | 305.15 (294.93 – 315.72) | 0.01 | -2.16 | .142 | 0.04 |
| **Block** | | | | | | |
| Block 2 | 1.08 (1.06 – 1.10) | 311.97 (302.65 – 321.58) | 9.20e-03 | 9.13 | **< .001** | 1.11 |
| Block 3 | 1.10 (1.08 – 1.12) | 316.78 (307.31 – 326.54) | 9.34e-03 | 10.93 | **< .001** | 1.61 |
| **Random effects** | *Coefficient* | | | | | |
| SD (Residual) | 0.07 | | | | | |
| SD (Intercept: Subject) | 0.08 | | | | | |
| SD (Intercept: Subject:Light) | 0.05 | | | | | |
| SD (Intercept: Subject:Block) | 9.25e-03 | | | | | |
| Adjusted / Unadjusted ICC | | 0.645 / 0.564 | | | | |
| Conditional / Marginal $R^2$ | | 0.690 / 0.125 | | | | |

**Table S8.** Linear Mixed-Effects Model analysis of mean reaction time on the Psychomotor Vigilance Task. Significant *p*-values corrected for multiple comparisons using Tukey's HSD are highlighted in bold. Cohen's partial $f^2$ was approximated as effect size[8]. BWL = Bright White Light. CI = Confidence Interval. ICC = Intra-Class Correlation Coefficient. LWL = Long-Wavelength Light. RT = Reaction Time. SD = Standard Deviation. SE = Standard Error. SWL = Short-Wavelength Light.

| | RTV | | | | | |
|---|---|---|---|---|---|---|
| *Parameters* | *Coefficient (95% CI)* | *EMM (95% CI)* | *SE* | *z* | $p_{corr}$ | $r_p$ |
| Intercept | 1.41e-08 (0.00 – 0.00) | | 2.37e-08 | -1.8.62 | | |
| Mean RT | 49.34 (35.34 – 68.90) | | 8.40 | 22.89 | **< .001*** | 0.96 |
| **Light** | | | | | | |
| SWL | 1.07 (0.99 – 1.16) | 66.59 (61.23 – 72.42) | 0.04 | 1.72 | .314 | 0.27 |
| LWL | 1.07 (0.99 – 1.16) | 66.38 (61.04 – 72.19) | 0.11 | 1.65 | .352 | 0.26 |
| BWL | 1.03 (0.95 – 1.11) | 63.74 (58.41 – 69.55) | 0.12 | 0.65 | .916 | 0.10 |
| **Block** | | | | | | |
| Block 2 | 0.87 (0.81 – 0.93) | 62.51 (57.65 – 67.78) | 0.03 | -3.84 | **< .001** | -0.52 |
| Block 3 | 0.83 (0.77 – 0.90) | 59.97 (55.28 – 65.06) | 0.03 | -4.82 | **< .001** | -0.61 |
| **Random effects** | *Coefficient* | | | | | |
| SD (Residual) | 0.26 | | | | | |
| SD (Intercept: Subject) | 0.20 | | | | | |
| SD (Intercept: Subject:Light) | 0.08 | | | | | |
| SD (Intercept: Subject:Block) | 0.08 | | | | | |
| Adjusted / Unadjusted ICC | | 0.438 / 0.156 | | | | |
| Conditional / Marginal $R^2$ | | 0.800 / 0.645 | | | | |

**Table S9.** Generalized Linear Mixed-Effects Model analysis of intra-individual reaction time variability on the Psychomotor Vigilance Task. Significant *p*-values corrected for multiple comparisons using Tukey's HSD are highlighted in bold. *p-value for the effect of mean RT is uncorrected as the variable is a continuous predictor. Pearson's partial *r* ($r_p$) was approximated as effect size[7]. BWL = Bright White Light. CI = Confidence Interval. EMM = Estimated Marginal Mean. ICC = Intra-Class Correlation Coefficient. LWL = Long-Wavelength Light. RT = Reaction Time. RTV = Reaction Time Variability. SD = Standard Deviation. SE = Standard Error. SWL = Short-Wavelength Light.

| | Mean 10% fastest RTs | | | | | |
|---|---|---|---|---|---|---|
| *Parameters* | *Coefficient (95% CI)* | *EMM (95% CI)* | *SE* | $t$(428) | $p_{corr}$ | $f^2$ |
| Intercept | 237.89 (231.90 – 244.04) | | 3.09 | 421.39 | | |
| **Light** | | | | | | |
| SWL | 0.98 (0.97 – 1.00) | 244.80 (238.77 – 250.98) | 7.82e-03 | -2.35 | .094 | 0.05 |
| LWL | 0.98 (0.96 – 0.99) | 239.36 (233.57 – 245.30) | 7.79e-03 | -2.82 | **.029** | 0.08 |
| BWL | 0.99 (0.97 – 1.00) | 241.13 (235.19 – 247.22) | 7.99e-03 | -1.86 | .252 | 0.03 |
| **Block** | | | | | | |
| Block 2 | 1.04 (1.03 – 1.05) | 244.24 (238.46 – 250.17) | 6.70e-03 | 6.28 | **< .001** | 0.53 |
| Block 3 | 1.05 (1.03 – 1.06) | 245.48 (239.66 – 251.43) | 6.73e-03 | 7.06 | **< .001** | 0.67 |
| **Random effects** | *Coefficient* | | | | | |
| SD (Residual) | 0.05 | | | | | |
| SD (Intercept: Subject) | 0.07 | | | | | |
| SD (Intercept: Subject:Light) | 0.02 | | | | | |
| SD (Intercept: Subject:Block) | 0.02 | | | | | |
| Adjusted / Unadjusted ICC | | 0.715 / 0.670 | | | | |
| Conditional / Marginal $R^2$ | | 0.732 / 0.062 | | | | |

**Table S10.** Linear Mixed-Effects Model analysis of the mean 10% fastest reaction times on the Psychomotor Vigilance Task. Significant $p$-values corrected for multiple comparisons using Tukey's HSD are highlighted in bold. Cohen's partial $f^2$ was approximated as effect size[8]. BWL = Bright White Light. CI = Confidence Interval. ICC = Intra-Class Correlation Coefficient. LWL = Long-Wavelength Light. RT = Reaction Time. SD = Standard Deviation. SE = Standard Error. SWL = Short-Wavelength Light.
